# Accurate detection of pathogenic structural variants guided by multi-platform comparison

**DOI:** 10.1101/2025.05.21.655285

**Authors:** Nico Alavi, M-Hossein Moeinzadeh, Jakob Hertzberg, Uira Souto Melo, Lion Ward Al Raei, Paolo Infantino, Maryam Ghareghani, Marco Savarese, Stefan Mundlos, Martin Vingron

## Abstract

Structural variants (SVs) are a common cause of human diseases and greatly contribute to inter-individual variability. Their detection represents a significant challenge due to their diversity, size, and enrichment in repetitive regions. With the use of high quality long-read technologies, the majority of these challenges can now be mitigated. However, many downstream applications ranging from clinical diagnostics and genome-wide association studies to large-scale aggregation of population variants continue to rely on short-read-sequencing data due to its high through-put and cost-effectiveness. Thus, the challenges of short-read sequencing SV detection remain a constant and relevant obstacle. We created *dicast*, a machine-learning method to improve the identification of true-positive SVs from short-read sequencing using genomic context and alignment features. *Dicast* is driven by a novel and comprehensive benchmark call-set created through the combination of several sequencing technologies and rigorous manual curation. This benchmark set served as the basis for a systematic evaluation of five sequencing platforms and fifteen SV detection methods across different SV classes, sizes, and genomic contexts, enabling us to quantify the strengths and weaknesses of each and inform the training of our model. Leveraging these insights, *dicast* outperforms state-of-the-art short-read-based callers and consensus approaches, identifying considerably more true-positive variants while maintaining a high precision. We also demonstrate the method’s applicability in diagnostic scenarios using putative pathogenic candidates in a limb malformation cohort, as well as known pathogenic variants in atrial fibrillation and neuromuscular disease cohorts. *Dicast* identifies all known pathogenic variants and 20% more manually confirmed candidate deletions than a consensus approach, and therefore can be used to reliably reduce time-consuming manual inspection in diagnostics.

**Graphical Abstract:** 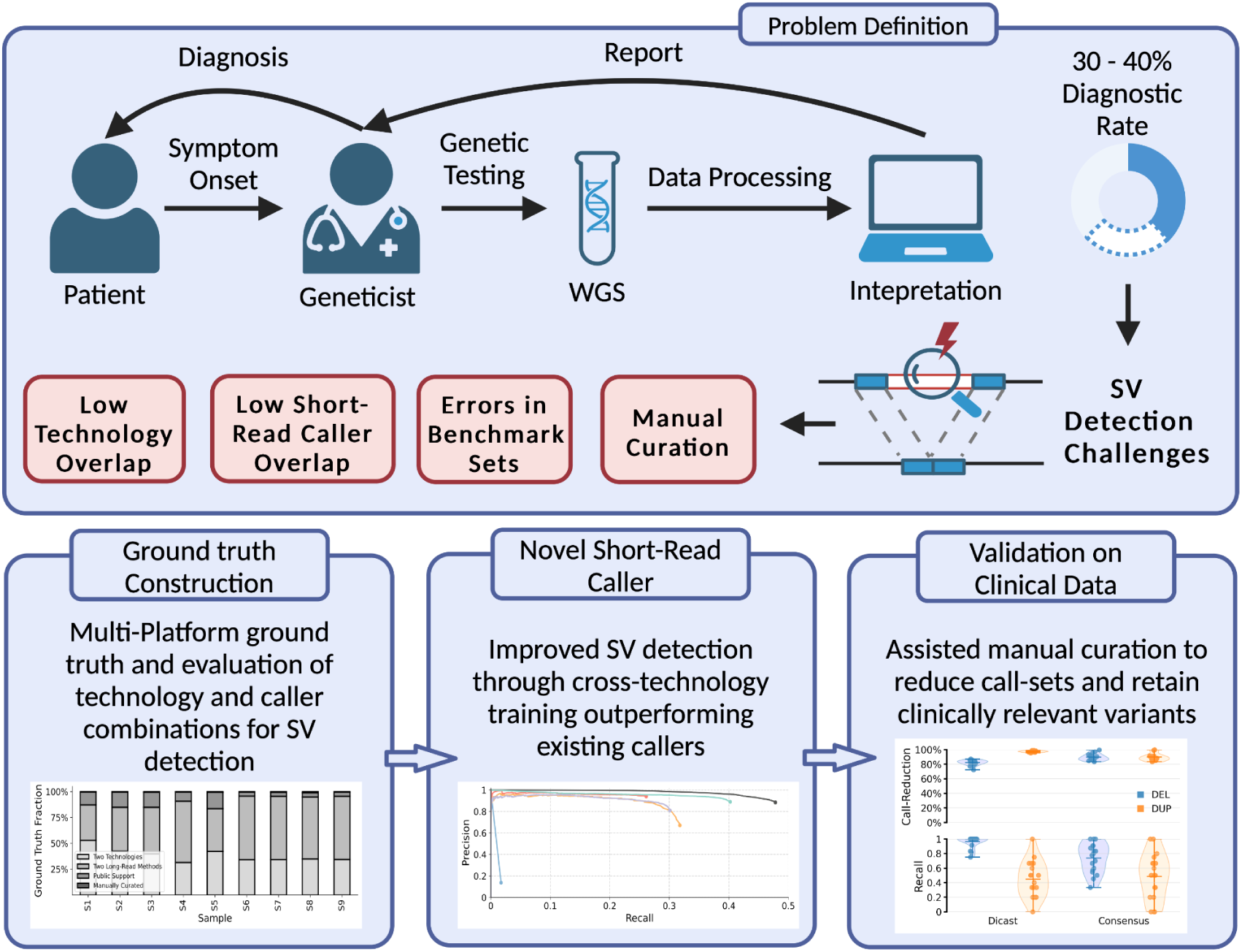

## Introduction

Despite significant advances in our understanding of the human genome, the diagnostic rate for genetic diseases remains low, leaving many patients without a molecular diagnosis (Taylor et al., 2015; Wortmann et al., 2015; Clark et al., 2018; Powis et al., 2018). Structural variants (SVs) contribute to a wide range of genetic disorders, yet their detection and interpretation remain challenging, limiting their integration into routine clinical diagnostics (Weischenfeldt et al., 2013; Liu et al., 2022). Improving SV detection, along with understanding the strengths and limitations of available sequencing technologies, is crucial for increasing diagnostic yield and identifying the genetic basis of previously unresolved cases.

When designing diagnostic pipelines, one of the first considerations is choosing the appropriate sequencing technology. Nowadays, there exists a myriad of technologies to read DNA. Short-read sequencing (SRS) typically generates paired-end reads with a size of around 100-150 bp (Buermans et al., 2014). During linked-read sequencing (LiRS), a short nucleotide-based barcode is attached to each read indicating which reads originate from the same DNA molecule, providing long-range information which can be utilized for phasing and the detection of SVs (Zheng et al., 2016; Chen et al., 2020). Long-read sequencing (LRS) generates reads ranging from several kilobases to over 100 kb (Rhoads et al., 2015). Additionally, optical genome mapping (OGM) constitutes an orthogonal technology to DNA sequencing where molecular labels are enzymatically added to specific motifs, and the labeled DNA is imaged to detect large structural variations by comparing label patterns to a reference genome (Yuan et al., 2020).

Alongside the choice of sequencing technology, researchers, and clinicians must select from an array of computational SV detection methods. Due to limitations in precision and recall of individual detection methods, it is common practice to apply multiple methods on the same patient data and use subsequent integration strategies such as consensus calling (Cameron et al., 2019). Consensus calling integrates results from multiple detection methods by filtering for variants detected by at least *n* out of *N* callers. A higher *n* improves precision by reducing false positives, while a lower *n* enhances recall by capturing more true variants at the cost of additional false positives. Due to this inherent tradeoff, better integration strategies are needed, in particular for short-read-based methods which exhibit high false-positive rates.

Accurately evaluating sequencing technologies, SV callers and integration strategies is non-trivial. Even though there exist several ground-truth SV datasets which these evaluations are based on, they can contain inaccuracies due to shared biases among technologies and methods. Previous efforts to assess SV detection performance were often confined to comparing individual SV callers without attempting to systematically assess the strengths and weaknesses of the underlying sequencing technologies (Zook et al., 2020, Ebert et al., 2021). Furthermore, Khayat et al. demonstrated the existence of notable biases in structural variation detection, particularly in how mapping methods, sequencing centers, and replicates contribute to analytical variability. Their findings highlight the need for multi-sample evaluations (Khayat et al., 2021). Additionally, in comparison to single nucleotide polymorphisms (SNPs), there is a lack of a definitive method or gold standard to determine the similarity between SVs. A robust evaluation requires the continuous improvement of existing SV benchmark datasets and the generation of novel ones.

Following detection and prioritization of variants, a key bottleneck in the diagnostic pipeline is the manual inspection of candidate SVs. This step ensures that each variant call is supported by sufficient sequencing evidence, reducing the risk of false positives. It usually relies on genome browsers such as *IGV* for visualizing read alignment at variant breakpoints, but quickly becomes time intensive as the number of potential variants grows (Robinson et al., 2011). Although tools like *SVHawkEye* and *Samplot* have been developed to facilitate this process, they either cater to specific sequencing technologies or do not provide sufficient information for comprehensive curation (Xiao et al., 2024; Belyeu et al., 2021).

In this work, we address these challenges by creating a ground truth SV database based on nine samples, leveraging multiple sequencing technologies to ensure a broad and accurate representation of SVs. To streamline the laborious process of manual curation, we introduce a novel technology-agnostic visualization tool, *cuban*, that enables scalable inspection of SV calls, ultimately supporting the development of our curated SV benchmark set. Using this benchmark, we systematically evaluated five different sequencing technologies and fifteen SV detection methods, assessing their strengths and weaknesses regarding SV type, size, and genomic context across multiple samples. We also explored the benefits of combining multiple technologies to gain a more comprehensive view of the structural variation present in a patient. Given that short-read sequencing remains the workhorse of routine clinical diagnostics, we then developed a novel machine learning model, *dicast*, which was trained on our ground truth database to learn sequencing patterns of true-positive SVs. *Dicast* outperformed all other tested short-read-based SV callers as well as consensus calling strategies in detecting insertions, deletions, and duplications. Finally, we assessed *dicast’s* clinical utility in three rare disease cohorts - congenital limb malformations, atrial fibrillation, and neuromuscular disorders.

## Results

### Establishing a High-Quality, Manually Curated Benchmark Set for SV Detection

In total, we sequenced nine patient samples employing a comprehensive set of sequencing technologies including Illumina SRS, PacBio LRS, two linked-read sequencing technologies by 10x Genomics and Universal Sequencing (TELL-Seq) as well as Bionano OGM (Fig. 1A). After preprocessing, we employed a total of 15 different SV detection methods, partly read-alignment based and partly *de novo* assembly based, to generate a comprehensive set of SV calls across our cohort (Fig. 1B). To generate a ground truth dataset and subsequently evaluate different methods and sequencing technologies, it needed to be determined which called variants were true positive (TP) variants and which were false positive (FP) variants.

**Fig. 1:**
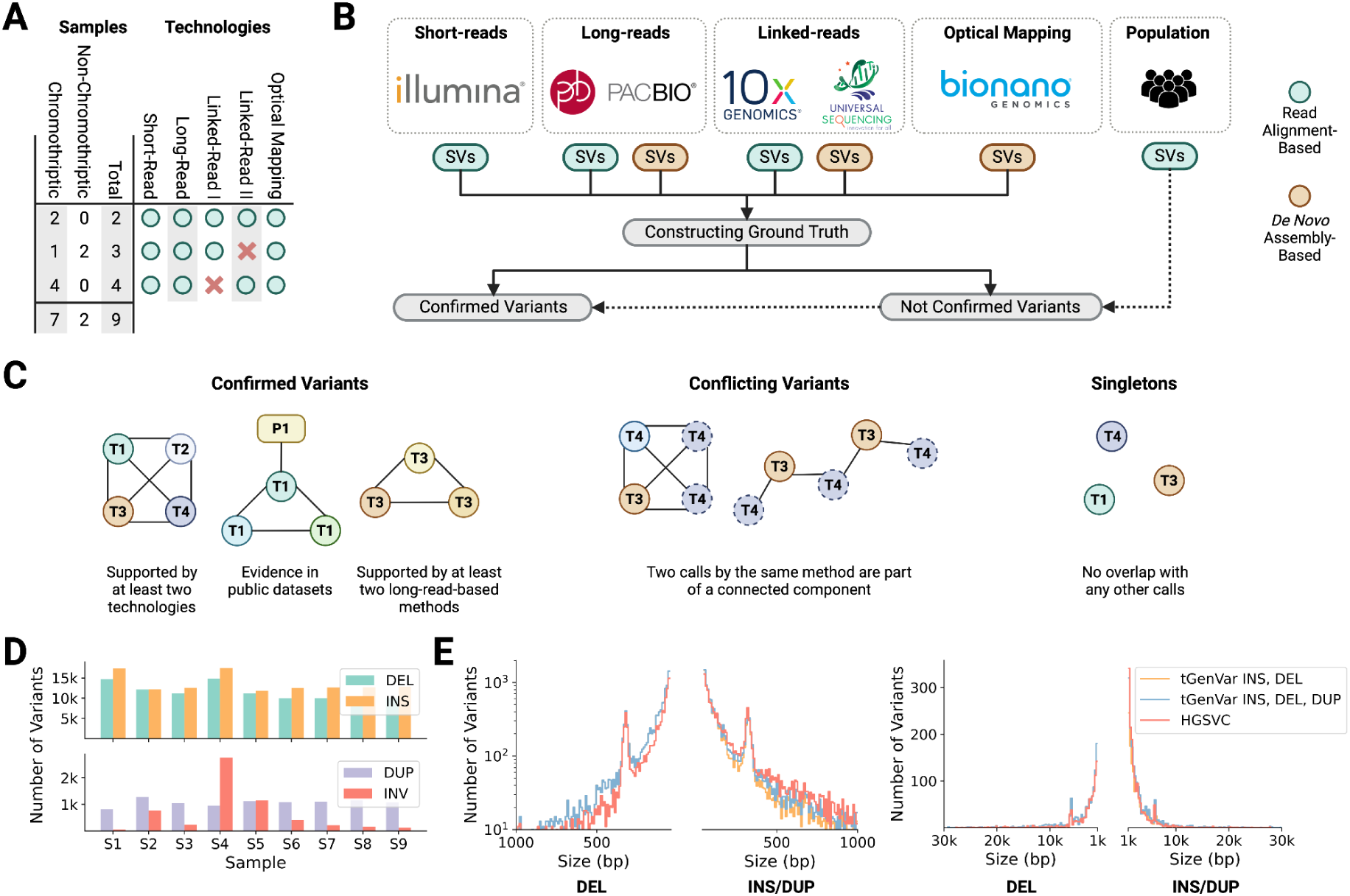
Establishing a High-Quality, Manually Curated Benchmark Set for SV Detection. A) Overview of samples in the ground truth and technologies they were sequenced with. B) Workflow for constructing the ground truth set of structural variant calls. C) Ground truth construction was performed using an overlap graph. Different types of graph substructures corresponding to confirmed variants, conflicting variants and singletons are shown. D) Number of variants in the ground truth dataset per sample, stratified by SV type. E) SV size distribution for SVs < 1kb (left) and between 1kb and 30 kb (right). The constructed ground truth set is being compared to the HGSVC Freeze 4.0 HG002 benchmark dataset.

For that, we have developed a novel graph-based framework. In this graph, individual SV calls are nodes which are connected via an edge if the breakpoints of two variants from different methods are at most 500 bp apart and their size similarity is at least 70%. Connected components in this graph correspond to structural variants which are supported by multiple methods. We call a variant confirmed, *i.e.*, a TP structural variant, if its respective connected component fulfills one of the following three criteria: 1) The variant is supported by at least two methods based on two different sequencing technologies, 2) the variant is supported by at least two long-read based methods, 3) the variant is supported by evidence from a publicly available database (see Fig. 1C). In contrast, we call variants conflicting if their connected component contains two nodes from the same method. After creating this first draft of a ground truth dataset, we systematically identified potential errors in our dataset. Potential FP or false negative (FN) variants were manually inspected by a panel of human curators utilizing a novel SV visualization tool, called *cuban*. Given the breakpoints of a candidate SV, *cuban* creates an image displaying different aspects of the aligned sequencing data to enable human curators to assess the variants’ validity. These aspects include coverage (compared to the chromosomal average), the presence of repetitive elements, distributions of insert size outliers and discordant read pairs as well as read alignments around the breakpoints (Sup. Fig. 12). We manually curated a total of 11,511 SVs, to mitigate potential biases arising from the automated confirmation process. This encompasses the removal of FP variants from the benchmark and the inclusion of previously excluded SVs.

The final ground truth SV callset consists of 122,133 insertions, 104,099 deletions, 9,664 duplications, and 5,877 inversions across all nine samples. On average, each sample possessed 13,570 insertions, 11,567 deletions, 1,074 duplications, and 653 inversions (Fig. 1D). Of the variants in the ground truth dataset 38% were on average supported by at least two methods from two different sequencing technologies, while 52% were included because they were detected by at least two long-read based methods. Furthermore, 9% were supported by public databases and 1% were added by identifying FN calls through manual curation (Sup. Fig. 2A,D). The majority of structural variants in our ground truth were supported by at least one long-read-based method (Sup. Fig. 2C).

We conducted a comparison between one sample from our newly constructed benchmark dataset - *tGenVar* - and the published HGSVC Freeze 4.0 benchmark dataset for HG002 (Ebert et al., 2021), focusing on the length distribution and composition of variants. Compared to our per sample average of approximately 13.5k insertions, 11.5k deletions, 1k duplications and 650 inversions, the HGSVC benchmark for HG002 contains 13.9k insertions, 9k deletions, 115 inversions and no separate set of duplication calls. In Figure 1E, we compared the size distribution of insertions and deletions, distinguishing between small SVs (<1 kb) and large SVs. The size distribution of deletions was similar between the two datasets, however our ground truth database exhibited a lower number of insertions. It is worth noting that both databases showed a peak around 350 bp for insertions and deletions, which can be attributed to the presence of Alu elements, a type of transposable element frequently found in the human genome. Additionally, a second peak observed around 1.5 kb is associated with Short Interspersed Nuclear Elements (SINEs).

### Comprehensive Evaluation of SV Detection Methods and Sequencing Technologies

Leveraging our newly constructed ground truth dataset, we systematically assessed each technology, method, and integration strategy for identifying structural variants across diverse sizes and genomic contexts, with the goal of providing a concise guide for clinicians and researchers. In total, we evaluated 15 SV detection methods across four sequencing and one optical mapping technology. We employed two different caller integration strategies to systematically evaluate the potential of each technology across varying SV sizes and genomic contexts. To ensure robustness, the benchmarking was performed across nine samples.

Among the long-read-based methods, *debreak* and *cutesv* consistently outperformed other tools by identifying the highest number of variants across multiple SV types. Specifically, *debreak* excelled in detecting insertions, deletions, and duplications, while *cutesv* showed particular strength in identifying insertions, deletions, and inversions (Fig. 2A,B; Sup. Fig. 3A,B). *Manta* was the most effective standalone short-read-based caller, though, like other SRS-based methods, it struggled particularly with detecting insertions (Fig. 2A,B). The use of barcoded linked-reads improved variant detection compared to conventional short-read methods but at the cost of lower precision, especially for duplications and inversions. Optical genome mapping demonstrated moderate effectiveness overall, but notably exceeded short-read-based approaches in insertion detection (Fig. 2A,B).

**Fig. 2:**
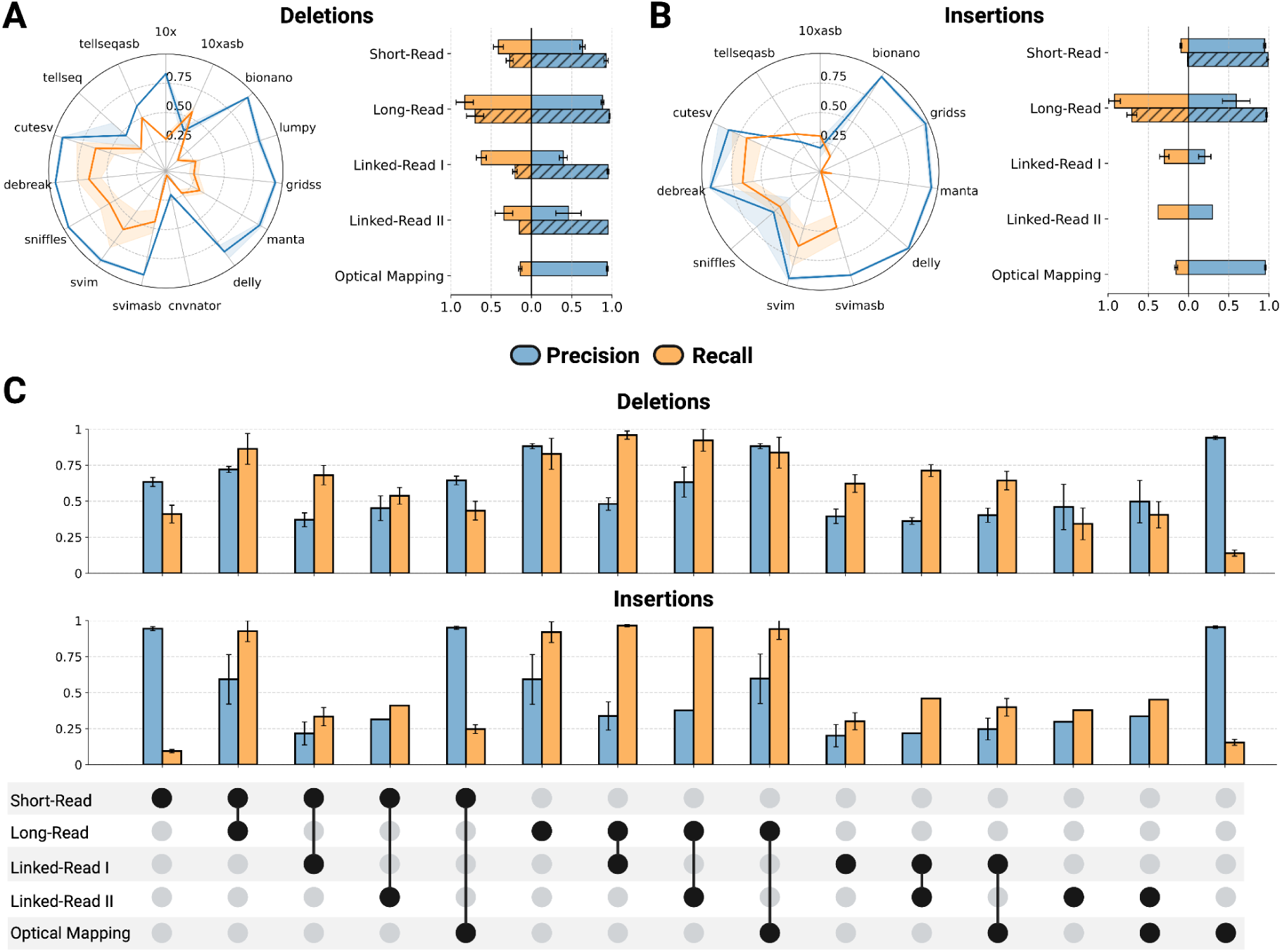
Comparative analysis of methods and sequencing technologies for structural variant detection. A) Precision and recall for deletion detection. The blue line represents precision, while the orange line represents recall. The confidence interval indicates the standard deviation for precision and recall across nine samples. The horizontal bar plots compare the precision and recall when combining multiple methods within one technology. Striped bars describe a more than 2/n caller strategy, while non-striped bars describe a union strategy (Short-Read: Illumina, Long-Read: PacBio CLR, Linked-Read I: 10x Genomics, Linked-Read II: TELL-Seq, Optical Mapping: Bionano) B) Precision and recall for insertion detection. C) Evaluation of combining multiple sequencing technologies for SV detection. Multiple methods within one technology were combined using the union strategy. Error bars indicate standard deviation across all samples.

To facilitate the comparison of sequencing technologies rather than just individual methods, we employed two caller integration strategies. Specifically, we used a union-based approach (combining variants from all methods based on a particular technology) as well as a more conservative consensus strategy (requiring agreement from multiple callers). The union strategy maximized recall but reduced precision, whereas the consensus approach increased precision significantly but at the expense of recall. Long-read sequencing demonstrated the highest TP detection rates for structural variants, consistently identifying a large proportion of variants, as shown in Figure 2A,B and Sup. Figure 3A,B. Precision remained high when combining results from multiple LRS-based callers. In comparison, SRS-based methods were notably less effective, detecting fewer than half of the benchmark deletions and fewer than 10% of insertions. While SRS precision could be increased substantially through consensus strategies, the overall recall was notably lower compared to LRS. The use of barcoded linked-reads, which allow for an additional de novo assembly step before SV detection, led to an increase in detected DELs and insertions but resulted in a decrease in precision compared to conventional SRS methods. Except for a negligible number of variants, linked-read-based methods failed to detect DUPs and INVs. OGM showed modest performance overall but demonstrated superior effectiveness to standard SRS methods in identifying insertions, with consistently high precision.

To identify the specific strengths and weaknesses of each sequencing technology in regard to SV detection, we assessed their performance in detecting variants across different size classes and genomic contexts. First, we divided all callsets as well as the benchmark sets into variants of sizes 50 bp - 100 bp (small), 100 bp - 1 kb (medium), 1 kb - 10 kb (large), > 10 kb (very large). SRS-based methods excelled at detecting very large deletions and small duplications but faced limitations detecting insertions larger than 1 kb due to read-length constraints. LRS-based methods demonstrated robust detection for variants up to 10 kb though precision declined for very large variants. Optical genome mapping was especially powerful for detecting very large variants, in particular, surpassing other technologies in detecting very large insertions (Sup. Fig. 3D).

In most cases, SV detection relies on the alignment of reads to the reference genome, which can vary in difficulty across different genomic regions. For example, the presence of repetitive sequences complicates the alignment process. Consequently, the ease or difficulty of SV detection varies depending on the specific genomic region, its repeat content and structure. Therefore, we investigated context-dependent SV detection performance for each evaluated technology. This was done by comparing the overall performance of a technology with its performance when constraining the evaluation to variants overlapping with certain genomic annotations (see Methods). LRS-based methods are the least impacted by repetitive genomic sequences, reflected by its stable recall across contexts (Sup. Fig. 3E). SRS and OGM-based methods display a drop of approximately 50% in recall in regions containing variable number tandem repeats (VNTRs) and CpG islands. In contrast, short and long interspersed nuclear elements (SINEs and LINEs), long terminal repeats (LTRs) and short tandem repeats (STRs) have a positive impact on their recall. Precision was most negatively impacted in short-read-based methods, with drops observed in CpG islands, low complexity regions, and retroposons.

Lastly, we evaluated potential synergistic effects on SV detection performance when combining methods from two different sequencing technologies. Overall, we did not observe major improvements in recall across all technology combinations and SV types (Fig 2C & Sup. Fig. 3C). The only modest improvement observed was when combining SRS- and OGM-based methods for detecting duplications.

### Machine learning reliably distinguishes true variants from artifacts

Despite the superior performance of long-read-based SV calling methods, short-read sequencing remains the most commonly used sequencing method in clinical diagnostics. This dominance is partly due to the lower cost of sequencing, availability of existing sequencing infrastructure, and scalability of the technology. To address the previously described shortcomings of SRS-based SV detection, we have developed a novel machine-learning-based SV detection method for deletions, insertions, and duplications termed *dicast*. *Dicast* obtains candidate SV calls either from unfiltered callsets produced by existing methods or from population catalogs. Each candidate SV gets annotated with more than 100 features, including the variant’s size and type, the alignment patterns of surrounding reads, and the genomic context the variant is located in. These annotations are then fed into an XGBoost model - trained on eight of the nine samples from our newly constructed benchmark - to distinguish true-positive SVs from false-positive artifacts.

To ensure comparability, we evaluated *dicast*, on an independent public benchmark dataset published by the Human Genome Structural Variation (HGSVC) consortium as well as on the one remaining sample of our newly developed benchmark. We called SVs using the short-read-based methods *delly*, *manta*, *lumpy*, *gridss*, and *cnvnator*. The resulting unfiltered callsets as well as a population catalog of variants served as input for *dicast* which subsequently predicted the probability of each variant to be present in the sample. For deletions and insertions, each callset was compared to a manually curated version of the HGSVC benchmark. Since at the time of writing, there does not exist a public benchmark dataset for duplications on GRChg38, we performed the evaluation based on one held-out sample from our newly constructed benchmark dataset.

*Dicast* outperformed all five tested SRS-based SV detection methods (Fig. 3A). For deletions, *dicast* achieves a best-in-class recall of 0.50 while maintaining a precision of 0.86. In contrast, the second-best short-read-based caller, *manta*, achieves a recall of 0.40 and a comparable precision of 0.89. Furthermore, we detect 40% of TP insertions compared to 10% detected by the second-best method, *manta*. For duplications, we achieve a recall of 0.27 and a precision of 0.89 using our current model. The second best evaluated short-read-based method, *gridss*, achieved a recall of 0.23 with a precision of 0.83. When population catalog variants were excluded as input to *dicast*, overall performance remained largely unchanged. However, recall for insertions declined noticeably, as most candidate variants in this category originated from population catalogs due to the limited capability of the input SV callers to provide candidate SVs (Sup. Fig. 4A).

**Fig. 3:**
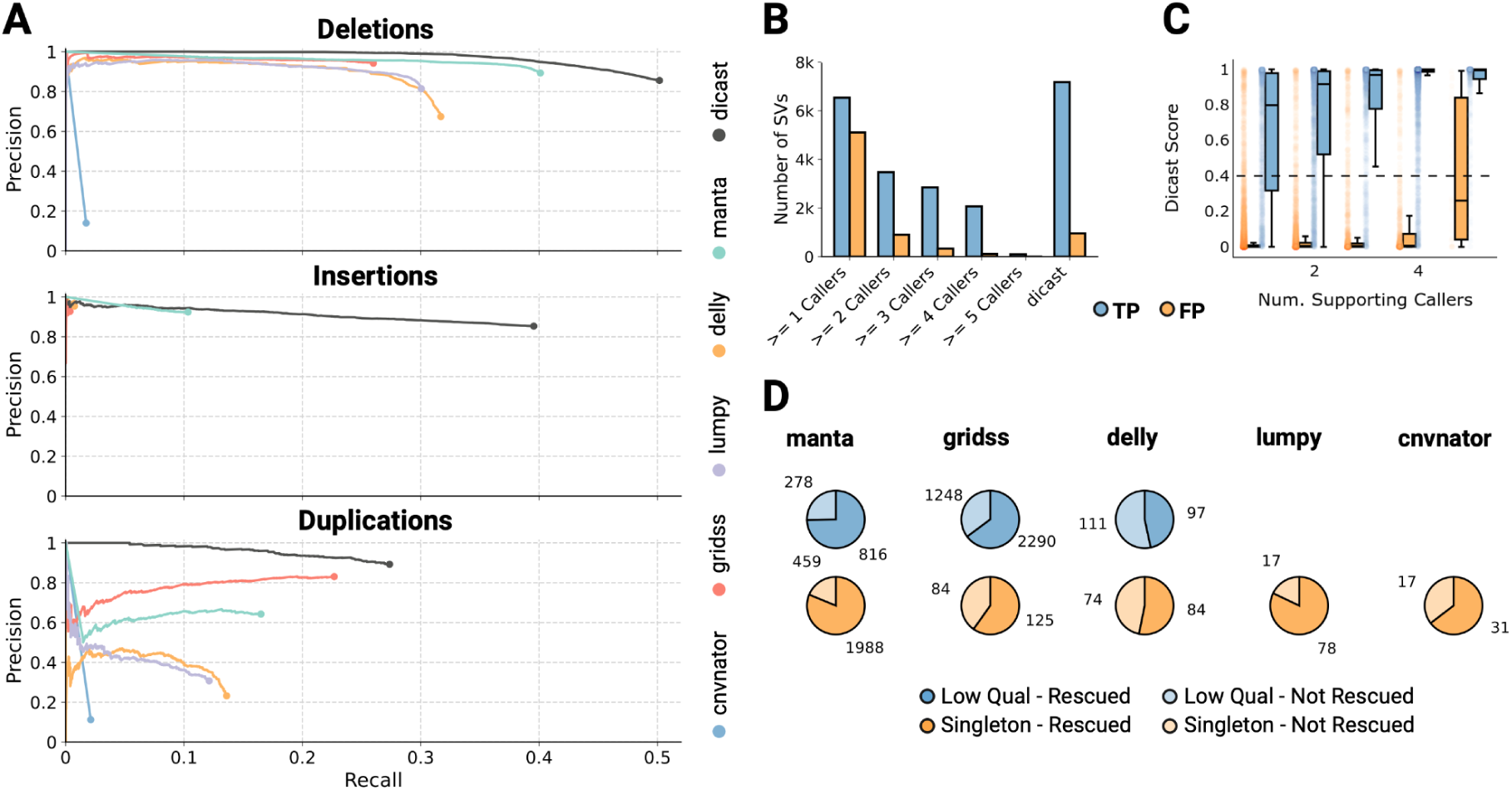
Novel machine learning tool reliably distinguishes true variants from artifacts. A) Precision-recall curves for detecting deletions, insertions, and duplications based on a manually curated version of the HGSVC Freeze 4.0 benchmark set for HG002 (DEL, INS) and one held-out sample from our newly constructed benchmark dataset (DUP). B) Number of TP and FP deletion and duplication calls when using *dicast* compared to various consensus calling strategies. C) Distribution of *dicast* scores depending on the number of input SV callers, a variant was detected by. D) Number of rescued variants, which are usually discarded in commonly employed SV detection pipelines. The first row describes the number of true variants which received a low-quality tag by the respective caller. The second row describes the number of true singletons, variants only detected by one SV caller, which would be filtered out using consensus approaches.

Next, we investigated the impact of a variant’s size on the detection performance for deletions and duplications (Sup. Fig. 4B). Across all size categories, *dicast* identified more or a comparable number of TP variants while maintaining an overall lower number of FPs. The largest improvement in terms of correctly identified variants can be observed for SVs between 50-500bp in size. For larger variants, the primary strength of *dicast* lies in filtering out FP artifacts while detecting a comparable number of TP SVs.

A commonly employed filtering strategy for structural variant calling workflows is consensus calling. Here, only variants detected by *n* out of *N* callers are kept, where *N* corresponds to the total number of callers being used and *n* to the caller support threshold. We compared various consensus calling strategies against using *dicast* on the unfiltered callsets of five SRS-based SV detection methods. *Dicast* identifies a comparable number of TP variants as when using the union of 5 short-read-based SV callers with 81% less FPs (Fig. 3B). When requiring a support of 2/5 callers, the consensus approach identifies fewer FPs SVs. However, this tradeoff comes with a cost of only detecting 52% of the TP variants identified by *dicast*. The ability of *dicast* in detecting TP variants only called by one or a few methods can be observed in Fig. 3C. While a higher number supporting SV callers corresponds to a higher median *dicast* score for TP SVs, the median score for SVs only detected by a single method is above 0.8 and above 0.9 for variants detected by two methods. Our methods’ ability to rescue commonly filtered out variants is further demonstrated in Fig. 3D. First, for each method, we calculated the number of TP variants which received a low-quality tag by their respective caller and would therefore be filtered out by commonly employed pipelines. *Dicast* assigns a majority of those variants (66%) a high quality score, thus rescuing them. Next, we calculated the number of TP variants which are only detected by an individual method, called singletons, and would therefore be filtered out in consensus calling workflows. *Dicast* rescued 78% of singleton deletions, insertions, and duplications.

In the medical field, only a select portion of a patient’s genomic variations have clinical importance. These important variations are typically those located near genes linked to the patient’s specific symptoms or condition. Wagner et al. have compiled a list of challenging medically relevant genes (CMRG) that are difficult to analyze due to their complex or repetitive nature (Wagner el al., 2022). Alongside, they have published a benchmark set of variants within these genes using sample HG002. We utilized this benchmark to evaluate the ability of commonly used SRS-based SV callers and *dicast* to detect deletions in CMRG. After filtering for deletions larger than 50 bp the benchmark contained 33 deletions. Out of those, *dicast* correctly identified 14 variants, one more than the best short-read-based method, *manta* (Sup. Fig. 4C). Currently, *dicast* is dependent on the candidate variants other methods provide. When providing *dicast* with a set of pre-defined regions to check, *dicast* correctly identified 30 out of 33 variants.

### Evaluating *Dicast*’s Potential to assist in Clinical Diagnostic Workflows

Manual Inspection of SVs is a critical step in clinical diagnostic workflows. It serves as the last *in silico* verification of potentially disease-causing genomic variants. Current standard workflows employ *IGV* or comparable applications visualizing aligned sequencing reads at breakpoint loci which are then inspected by a qualified clinician or bioinformatician. This practice becomes increasingly challenging with the high throughput from current sequencing technologies - especially when shifting from WES to WGS with only 1.8% of SVs and InDels occurring in Refseq coding exons (Huddleston et al., 2017). *Dicast* has the potential to automate this process or at least significantly reduce the number of candidate SVs that are to be inspected. Current diagnostics workflows already employ comparable strategies to minimize the number of detected candidate variants before manual curation, such as caller-specific filters and consensus calling (Koboldt et al. 2020). While these strategies can help to reduce the number of false-positives, they are often too rigorous and potentially exclude variants that prove to be clinically relevant. With *dicast* we aim to optimize this precision-recall trade-off and retain candidate SVs that would be discarded in current diagnostic workflows.

To evaluate *dicast* in the setting of clinical diagnostics, we implemented three pipelines for the detection and prioritization of putative pathogenic deletions and duplications centered around a cohort of 18 patients with limb malformations: a short-read pipeline representing the current standard for SV detection that relies on caller defined quality filters, a second short-read pipeline that includes all low-quality calls, and a pipeline for a short- and long-reads with caller defined quality filters. In each pipeline, we first reduced the initial call-set to rare variants. We then assessed their potential regulatory impact using a comprehensive set of disease-specific annotations, resulting in a set of functionally relevant SVs. A detailed description of this process is provided in the methods. Briefly, we processed complementary transcriptomic (RNA-seq) and chromatin conformation (Hi-C) data-sets for each patient. We then incorporated the processed data in a previously published annotation method and derived overlap-based features for rare SVs to guide the identification of functionally relevant calls (Hertzberg et. al, 2022). Finally, we identified TP candidates through extensive manual curation. These candidates serve as ground truth for our analysis of *dicast*’s performance in clinical diagnostic workflows.

We set out to evaluate *dicast* for three potential applications: 1) Reducing the number of regulatory variants that need to be manually inspected (“callset reduction”), 2) identifying (“rescuing”) low-quality candidates and 3) highlighting novel candidates otherwise only confirmed through additional long-read evidence. To evaluate *dicast* with regard to the first application, we applied the method to all functionally relevant calls identified with our second pipeline and assessed its performance based on the ground truth of manually curated candidates (Fig. 4B). *Dicast* is able to identify the vast majority (96.21%) of manually confirmed deletions as true-positives. Of the corresponding assigned probabilities, 87.50% are higher than the 95th percentile. We then compared *dicast’s* performance to a consensus calling approach that considers variants supported by more than one caller as true-positives (Fig. 4C). For deletions *dicast* clearly outperforms the consensus approach with the average recall of 0.96 compared to 0.74 and only a minor decrease in call-set reduction (82.23% vs. 89.65%) resulting in a median of 29 variants per sample that need to be manually curated. For duplications, both *dicast’s* and the consensus approaches recall are notably lower (0.45 and 0.48, respectively). However, in this case, *dicast* would achieve this recall with a considerably higher call-set reduction (97.51% vs. 89.85%). To evaluate *dicast* regarding the second potential application, i.e., rescuing low-quality calls, we applied the model to the subset of low-quality yet manually confirmed candidates from our second pipeline that would be discarded by caller specific filters (31 out of 230). *Dicast* was able to identify 12 out of 31 using a threshold of 0.4 while only a single low quality was supported by the consensus approach. Finally, for the third application, we investigated the proportion of confirmed candidates that would otherwise require the support of a second sequencing technology, i.e., those uniquely shared between the second and third pipeline. In this subset, *dicast* can identify all manually confirmed variants as true-positives.

**Fig. 4:**
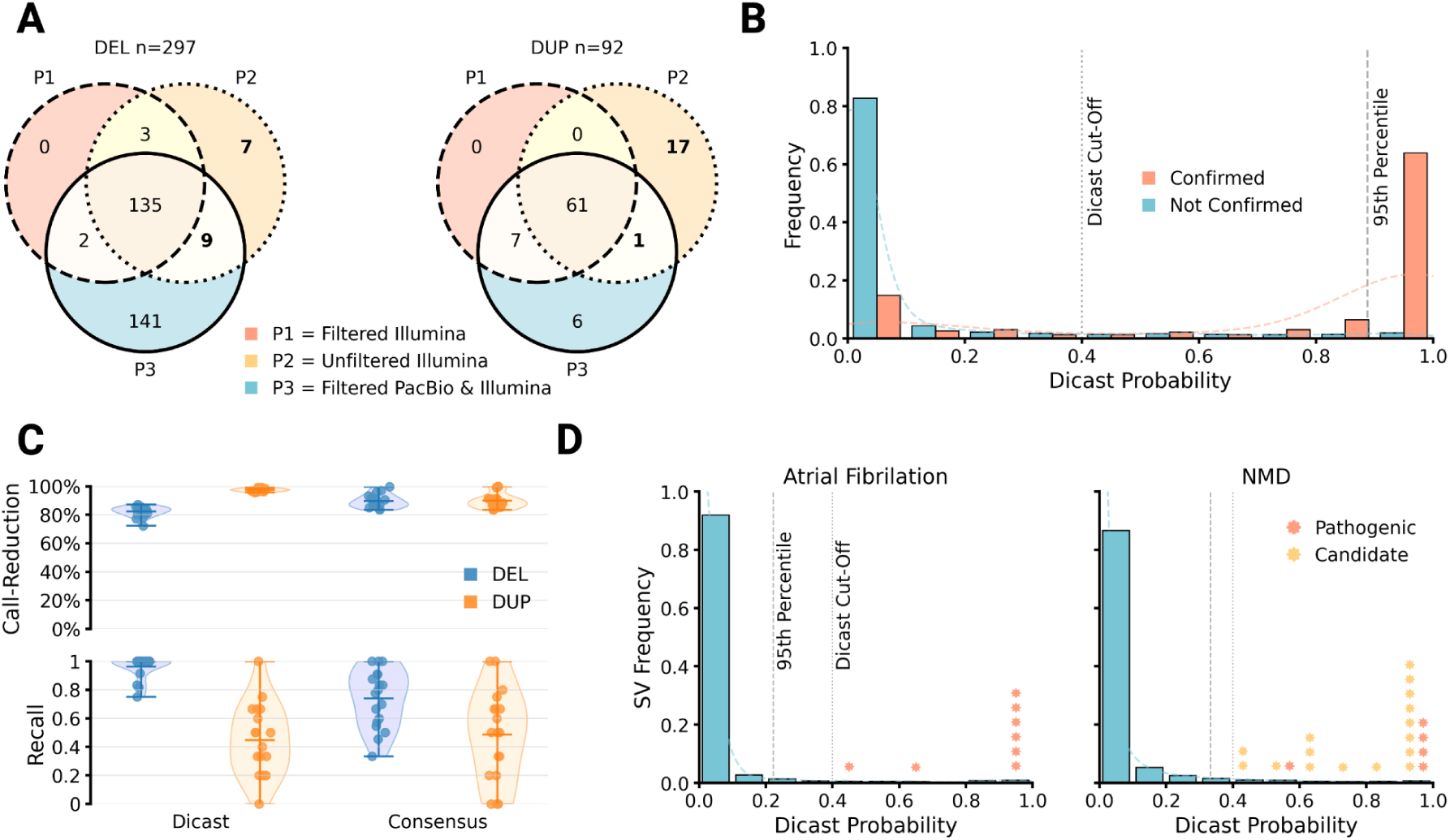
Evaluating the potential of *dicast* in the content of clinical diagnostic workflows. A) Overlap of manually curated candidates. The figure shows the overlap between the final manually curated sets (“Candidates”) of three pipelines for clinical diagnostics as an unweighted Venn diagram. B) *Dicast* Predictions for Candidate Variants. Here we show the frequency of *dicast* scores for functionally relevant calls grouped as manually confirmed candidates and variants discarded during manual curation. C) *Dicast* and Consensus performance. This figure compares the reduction in call-set size and recall based on the manually curated candidates when using Dicast or a Consensus approach. Dicast’s recall and call-set reduction for each sample is based on the recommended score threshold of 0.4. D) Dicast predictions for Disease Cohorts. In this figure, we display the frequency of *dicast* scores for functionally relevant deletion and duplications in two rare disease cohorts: Patients with atrial fibrillation (left subfigure) and neuromuscular diseases (NMD) (right subfigure). The absolute number of known pathogenic variants and candidates per *dicast* score bin are shown as red and orange markers, respectively.

The candidate call-sets generated using our three pipelines suggest that there is potential in considering low-quality calls in diagnostic workflows. However, harnessing that potential requires manual curation of approximately 5-times as many regulatory variants. This clearly highlights the need for methods like *dicast* to automate this process and maximize the recall of true candidate variants while minimizing the number of variants to be inspected.

Even though the ground truth set of manually curated variants are certain to be present in the affected individuals, their functional impact is yet to be confirmed. To evaluate *dicast’s* potential on known pathogenic variants (Sup. Table S1) in rare disease cohorts, we applied our short-read pipeline and *dicast* to an atrial fibrillation (AF) and a neuromuscular disease (NMD) cohort (Fig. 4D). We also employed the *Lucid Genome Suite (LGS)*, a platform for a collaborative analysis, for 20 previously unsolved NMD cases (Lucid Genomics, 2025). Via this visualization and interpretation framework, we collected experts’ opinions on putative pathogenic candidate variants, which were then subjected to downstream validation processes and included in this evaluation of *dicast*. In both cohorts, *dicast* can significantly reduce the number of functionally relevant calls with a call-set reduction of 96.70% (AF) and 96.01% (NMD) and correctly classifies all known pathogenic variants as true-positives.

## Discussion

The diagnostic journey for rare disease patients often extends over several years and requires substantial resources before achieving a definitive molecular diagnosis. Even with the dramatic advancements in genetic testing technologies, the current success rates are between 30-40% depending on the disease. This success rate is influenced by several critical factors, including the selection of sequencing technology and the downstream analysis of sequencing data encompassing variant detection, prioritization, and interpretation. The data analysis can further be split based on the investigated variant type. Single nucleotide variants have been studied extensively and are routinely assessed in clinical diagnostic workflows. The role of larger genomic alterations, however, is likely underestimated due to the challenges associated with their detection and interpretation.

In our analysis, we therefore focused on SVs in the context of clinical diagnostics, with the aim to overcome current bottlenecks during variant detection and consequently increase the potential for a successful diagnosis. We first created a ground truth database of structural variants consisting of nine samples and five different sequencing technologies. We utilized our newly constructed ground truth to conduct a comprehensive comparison of sequencing technologies, gaining insights into which technologies can be used to maximize SV detection sensitivity. Given the continued use of short-read sequencing due to its scalability and affordability, we developed *dicast* for short-read-based SV detection outperforming existing SV callers and consensus calling approaches. Finally, we successfully evaluated our model in several clinical scenarios.

The present work is subject to several limitations. The two-technology criterion used to include variants in our ground truth dataset does not account for the individual strengths of specific sequencing technologies. To address this, we also incorporated variants supported by at least two long-read-based methods or those common in the general population. However, this approach may have introduced false positive SVs into the dataset due to shared biases among different detection methods. While we made efforts to mitigate these biases by systematically manually inspecting potential FP and FN variants, this was only feasible for a subset of the final ground truth dataset. More extensive collaborative and distributed efforts in manual curation, e.g., through a public web application, will be crucial to improving the quality of SV truth sets in the future. Furthermore, when evaluating different SV detection methods, we focused exclusively on the presence or absence of variants, without considering their predicted genotypes. However, in clinical settings, the zygosity of a variant is a critical factor. Currently, *dicast* relies on candidate variants detected by other SV detection methods, limiting its maximum recall to the union of all input callers. To address this, we plan to develop new strategies to expand the set of candidate variants that *dicast* can evaluate, while also incorporating additional SV types beyond deletions, insertions, and duplications.

With *dicast* we take an important step towards a more accurate and comprehensive SV detection for short-read data. This has wide-ranging implications for multiple downstream applications. Here, we focused primarily on *dicast* benefits for clinical diagnostics, significantly reducing the number of false positives and the time-consuming manual curation of candidate variants allowing a rapid and confident evaluation of putative pathogenic variants. *Dicast* has, however, the potential to be used for other applications such as genome-wide-association studies (GWAS) and large-scale aggregation of population variants.

## Methods

### Sequencing and Optical Mapping

Illumina paired-end and PacBio CLR sequencing was performed as described in (Schöpflin et al., 2022). Linked-read sequencing by 10x Genomics and Universal Sequencing (TELL-Seq) as well as optical mapping by Bionano Genomics was performed by the respective machine manufacturer according to their best practices.

### Data Preparation and Structural Variant Detection

#### Illumina Paired-End WGS

Reads were aligned to GRCh38 using *bwa-mem* (Version 0.7.17, Li, 2013). The resulting *BAM* files were sorted, and duplicate reads were marked using *samtools sort* and *samtools mdup* (Version 1.19.2, Danecek et al., 2021), respectively. Structural variant detection was performed using five different SV callers: *delly* (Version 1.1.6, Rausch et al., 2012), *lumpy-smoove* (Version 0.2.8, Layer et al., 2014), *manta* (Version 1.6.0, Chen et al., 2016), *gridss* (Version 2.13.2, Cameron et al., 2021), and *cnvnator* with bin size 1kb (Version 0.4.1, Abyzov et al., 2011). All tools were run with default parameters, except indicated otherwise. HLA, decoy or alternate contigs and regions with much higher coverage than expected were excluded from *delly* and *lumpy* SV calling (https://github.com/hall-lab/speedseq/blob/master/annotations/exclude.cnvnator_100bp.GRCh38.20170403.bed).

#### PacBio CLR WGS

For each sample, subreads in *uBAM* format were pooled and converted to *FASTA* format using the tool *bam2fastq* (Version 1.3.0, Dexheimer et al., 2014). Pooled subreads were aligned to GRCh38 using *minimap2* (Version 2.15, parameters *-x map-pb -L -MD, Li, 2018*). Structural variants were called using the tools *svim* (Version 1.2.0, Heller et al., 2019), *sniffles2* (Version 2.0.7, Smolka et al., 2024), *cutesv* (Version 2.0.2, Jiang et al., 2020), and *debreak* (Version 1.0.2, Chen et al., 2023). All tools were executed with default parameters unless stated otherwise. For a subset of the samples, pooled subreads were *de novo* assembled with *FalconUnzip* (Version 0.5, Chin et al., 2016). The assembled diploid contigs were aligned to GRCh38 using *minimap2* (Version 2.15, parameters *–paf-no-hit -x asm5 –cs –r2k*). Finally, SVs were called using the tool *svim-asm* (Version 1.0.2, Heller et al., 2020) with default parameters.

#### 10X Genomics Linked-Read WGS

Linked-read *FASTQ* files generated by 10x Genomics were processed with *longranger* (Version 2.2.2, Marks et al., 2019). *Longranger* performs sequencing quality control, alignment to GRCh38, as well as detection of SNVs and SVs. For a subset of samples, *de novo* assembly has been performed using *supernova* (Version 1.2.0, Weisenfeld et al., 2017). Similarly to the long-read data, assembled contigs were aligned to GRCh38 using *minimap2* with the same command and structural variants were detected using *svim-asm*.

#### Universal Sequencing (TELL-Seq) Linked-Read WGS

To process TELL-Seq sequencing data with the longranger pipeline, the barcodes present in the FASTQ files needed to be converted to a longranger compatible format. For that, a tool called ust10x, provided by the sequencing provider Universal Sequencing, has been employed (Universal Sequencing, 2019). The tool converts the 18bp TELL-Seq barcode to a 16bp 10x compatible barcode while considering a whitelist of 10x compatible barcodes provided by the user. Subsequently, TELL-Seq FASTQ files were processed analogously to 10x Genomics FASTQ files using the longranger pipeline, *supernova* and *svim-asm*.

#### Bionano Optical Mapping

Raw data was processed using the software suite *Bionano Solve* (Version 1.5.3, Bionano Genomics, 2020). The output obtained from the software consisted of assembled consensus optical maps in *CMAP* format, aligned consensus maps to a reference genome in *XMAP* format, as well as structural variant calls in *SMAP* format. Unless stated otherwise, all subsequent tools and scripts used to further process Bionano data are part of the *Solve* software suite. *SMAP* files were converted to *VCF* format using the script *smap_to_vcf_v2.py* (Bionano Genomics, 2020). The format of the resulting *VCF* header was adapted using a custom Bash script. In particular, the file format specification line was moved to the correct position within the file and the length of the reference genome contigs was added to the respective header lines. Next, variants whose start position exceeded their end position were removed and variants were sorted according to their position using *bcftools sort* (Version 1.21, Danecek et al., 2021). Variant representations were standardized, and duplicate records removed using *bcftools norm* (Version 1.21, parameters -d all).

#### Population Databases

As part of the ground truth construction, SV calls from the analyzed samples have been compared to SV calls from publicly available databases. For that, we utilized data from multiple sources, including from the Genome in a Bottle (GIAB) SV Tier 1 v0.6, gnomAD SV v2.1, and Human Genome Structural Variation (HGSVC) Freeze 3 dataset (Zook et al., 2020; Karczewski et al., 2020; Logson et al., 2024). Additionally, we included SV calls from a population catalog published by Audano et al. (Audano et al., 2019). Variant calls which were only available on GRCh37 were lifted over to GRCh38 using the tool *liftover* (Genovese et al., 2024).

### Variant Quality Filtering

For comparing different sequencing technologies and SV detection methods, a ground truth set of variants needed to be established for each sample. First, a custom Python script was employed to convert *VCF* files across all methods to a standardized format. Subsequently, all *VCF* files were filtered to remove low-quality variants as defined by the respective callers. For the methods *delly*, *manta*, *gridss*, *cnvnator*, *svim*, *sniffles2*, *cuteSV*, *debreak* all variants which have been assigned the *PASS* tag were kept. Additionally, variants which have been assigned the *IMPRECISE* tag were removed. For *lumpy,* all variants with a quality score below 30 have been removed. The *longranger* pipeline, which was applied to 10x Genomics and TELL-Seq data, generated two structural variant *VCF* files, one for deletions and one for large SVs, per sample. For both files, variants with the filter tag *PASS* were kept and those with the tag *IMPRECISE* were removed. Both files were merged and missing reference contig lengths were added to the *VCF* header. For the converted optical mapping *VCF* files, variants whose start position exceeded their end position were removed, and variant representations were standardized, using *bcftools norm* (Version 1.21). Lastly, duplicate entries defined by SV type, chromosome, start position, and end position were removed using a custom Python script and *VCF* files were sorted using *bcftools sort*.

### Construction of Ground Truth

The construction of the ground truth can be divided into two major steps. First, it needed to be determined which variants in the filtered *VCF* files were indeed true-positive variants. Subsequently, true-positive variants needed to be merged to create a single SV ground truth for each sample. To identify true-positive SVs we employed multiple criteria, most of which are based on an initial pairwise overlap between all structural variants. For that, we employed the tool *truvari* (Version 2.2.0, parameters bench -p 0 –sizemin 10 –sizefilt 10 –sizemax 100000000 –dup-to-ins, English et al., 2022). The default parameters of *truvari* determine two SVs to be the same if their respective breakpoints are within 500 bps, and they exhibit a size similarity of at least 0.7. Because the nature of optical mapping does not allow for precise pinpointing of SV breakpoints, a different strategy was required to compare OGM *VCF* files to those generated by other methods. To implement this, each OGM *VCF* file was divided into 40 separate files, with each file representing a distinct quantile of the breakpoint confidence interval reported by Bionano’s *Solve* software. The region in which a breakpoint may lie is indicated by this confidence interval. Instead of performing a single comparison of the entire OGM *VCF* against all other *VCF* files, 40 pairwise comparisons were carried out—one for each quantile-based OGM *VCF* file. In each comparison, *truvari*’s –refdist and –chunksize parameters were set according to the relevant quantile. The resulting overlap sets were then merged in an iterative manner: overlaps involving OGM SVs with narrower confidence intervals were included first, and if the same variant from another method overlapped a higher-interval OGM SV, that subsequent overlap was disregarded.

This resulted in a sample-specific list of pairwise overlapping SV calls between all methods and public databases. Next, for each sample, a graph *G = (V, E)* was constructed, where each SV was represented as a vertex *v* ∈ *V*, and an edge *e* ∈ *E* indicated an overlap between two variants. Finally, connected components *C* ⊆ *V* within *G* were identified, where each connected component *C* is defined as a maximal set of vertices such that every pair of vertices in *C* is connected by a path in *G*. Here, connected components represent SVs which have been detected by multiple methods, each represented by a vertex *v* ∈ *C*. Several criteria were employed to select connected components, i.e., variant representations, to be included in the final benchmark set of SVs. First, variants which were supported by at least two methods based on two different sequencing technologies were selected. Furthermore, variants which were supported by at least two different long-read-based methods were included. Lastly, variants which were supported by multiple methods of the same technology and were present in any population database were included in the benchmark. This strategy resulted in a set of connected components consisting of multiple variant calls representing the same benchmark SV. Consensus start and end positions were inferred by calculating the sum of absolute differences between each candidate breakpoint and all other breakpoints, selecting the breakpoint pair (left and right) that minimized the total divergence from all other calls. Consensus sizes for insertions were inferred by selecting the most frequently occurring size when a value appeared multiple times in the calls, or when no consensus existed, by choosing the size that minimized the total absolute difference to all other size estimates.

### Manual Curation and Variant Visualization

Benchmark SV datasets, comprised of presumed true variants in an individual genome, can contain inaccuracies that arise from the methodologies used in their construction. These errors typically fall into two categories: FPs, which are erroneously included due to shared biases across technologies, and FNs, which are true variants missed because the strengths of certain technologies are underrepresented by multi-technology criteria. To mitigate such errors in our constructed ground truth datasets, we conducted extensive manual curation on a carefully selected subset of variants. We applied the following criteria to identify candidate variants for manual review:

- False Positives: Variants present in the benchmark with a dicast score < 0.4 were selected. Additionally, for each variant-calling method, we identified the 0.2 percentile of its internal quality score distribution and selected the 50 variants with the lowest scores.
- False Negatives: Variants absent from the benchmark but with a dicast score > 0.4 were selected. Similarly, for each method, we determined the 0.8 percentile of quality scores and selected the 50 highest-scoring variants.

Since dicast receives as input variants from the five used SRS-based methods, all variants came from these methods, not only benefitting dicast but also the method used to call this variant originally. The selected variants were manually reviewed by a panel of five human experts using a custom web application. This tool presented visual representations of each variant to facilitate evidence-based decisions on their validity. Based on expert consensus, FP variants were removed from the benchmark set, and FN variants were incorporated. To support manual curation, we developed a Python-based visualization tool named *cuban*. The tool generates PNGs that summarize key features of each SV, providing comprehensive context for expert evaluation (see Sup. Fig. 12 for a visual explanation). Information from the alignment of sequencing reads is extracted using *pysam* (Danecek et al., 2021). The visualizations include the following tracks:

#### Repetitive Elements

Repetitive sequences in the human genome can affect read alignment and variant detection. This track displays annotated repeat elements that are near the structural variant. VNTR and STR annotations were obtained from Lu et al., 2021. Other repeat element annotations were obtained from the *repeatmasker* track of the UCSC Table Browser (Karolchik et al., 2004).

#### Coverage

Read coverage across the whole structural variant with padding on both sides. Breakpoints are indicated by vertical black dashed lines. Average chromosomal coverage is indicated by a horizontal red dashed line and was calculated by randomly sampling the average coverage of 1,000 regions of size 1kb. For the coverage calculation, all reads are considered. A red coverage line displays the coverage when only considering reads with a mapping quality > 20.

#### Insert Size Outlier distribution

Insert size is the distance between two mates of a read-pair. A higher insert size of read pairs around the breakpoints than expected from the sequencing experiment can indicate the presence of an SV. The peaks of the line plot across the entire length of an SV display the number of reads that have an unusually high insert size (larger than 1 kb).

#### Discordant Read Pairs

The distribution of discordant read pairs is assessed across the whole length of the variant. Counts of read pairs in reverse-reverse (RR, orange), reverse-forward (RF, dark blue), forward-forward (FF, green) are displayed. Additionally, read counts where the two mates align to different chromosomes are shown (TX, red).

#### Read Alignments

In addition to the tracks across the entire length of the SV, *cuban* visualizes the alignment of reads within windows of 200 bp around each breakpoint. For that, aligned reads are collected with *pysam* and their CIGAR strings are fed into a matrix where each row corresponds to one read and each column to one nucleotide in the reference genome. The numbers in each cell correspond to different CIGAR string operations, which are subsequently visualized using the following color code:

- Match/ Mismatch: Light gray
- Low MAPQ: Dark gray
- Deletion: Blue
- Insertion: Yellow
- Soft-clip: Green
- Hard-clip: Red
- Split-read: Black stripes
- RR orientation: Orange stripes
- RF orientation: Dark blue stripes
- FF orientation: Green stripes

#### Read Connections

Split reads contain two segments from the same read that align to different locations in the reference genome. When these segments map to opposite sides of an SV, they can signal the presence of that SV. In such cases, the two aligned segments are connected by a black line in the visualization. Similarly, when two mates align to opposite sides of an SV, they are connected by a red line in the alignment view.

### Benchmarking SV Detection Methods and Technologies

#### Evaluation of SV Detection Methods

Based on the newly constructed ground truth dataset, a systematic evaluation of SV detection methods and their underlying sequencing or optical mapping technology was performed. A variant was considered confirmed if it overlapped with a benchmark variant using the following criteria: breakpoints within 500bp of each other and length similarity of at least 70% (for insertions, only breakpoint distance within 200bp was considered). Overlaps were performed using the Python library *bioframe* (Open2C et al., 2024). Performance metrics were then calculated by treating confirmed variants as true positives and unconfirmed variants as false positives, while benchmark variants without matching calls were counted as false negatives. Multiple calls which overlapped with the same benchmark variant were counted as a single TP variant, with the highest quality score being retained among the overlapping calls. Precision and recall were calculated as:

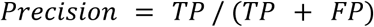

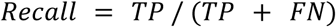

Evaluation results for each SV caller across multiple samples were aggregated by calculating mean and standard deviation.

#### Evaluation of Technologies

As a guide for choosing the right sequencing technology for various use cases, we aimed to not only compare SV detection methods but also the sequencing and optical mapping technologies these are based on. To facilitate this comparison, two caller integration strategies per technology were employed. First, a union call set was generated for each technology per sample by pooling all variants called by any of the associated callers. Performance (precision and recall) was then calculated for this union set against the benchmark dataset using the TP, FP, and FN definitions described above. Second, a consensus call set was generated for each technology per sample by retaining only those SVs called by at least 2 different SV detection methods. Again, performance metrics were assessed for this callset against the benchmark. Similar to the evaluation procedure of SV detection methods, performance metrics for each technology were aggregated across multiple samples by calculating mean and standard deviations.

Next, for each technology and SV type, performance was assessed for different size categories. The consensus callsets for each technology served as the basis for this analysis. Call and benchmark sets were divided into small (50-100 bp), medium (100 bp-1 kb), large (1-10 kb), as well as very large (> 10 kb) variants and precision and recall were computed as described above.

To assess the effect of genomic context on detection performance, the overall performance of a technology (defined by its consensus call set) was compared to its performance when constraining the evaluation to variants overlapping with certain genomic annotations. Genomic annotations used for this analysis were CpG islands, long interspersed nuclear elements (LINEs), short interspersed nuclear elements (SINEs), long terminal repeats (LTRs), low complexity regions, retroposons, simple repeats, short tandem repeats (STRs) as well as variable number tandem repeats (VNTRs). VNTR and STR annotations were obtained from a catalog published by Lu et al. (Lu et al., 2021) whereas CpG islands were downloaded from the UCSC Table Browser (Karolchik et al., 2004). All other annotations were extracted from the repeatmasker track of the UCSC Table Browser. Overlaps between annotations and variants were calculated using the Python library *bioframe*. A variant was assigned to an annotation if their corresponding intervals overlapped.

To assess the performance of different technology combinations, the union of their respective consensus call sets was formed. This served as the basis to calculate the number of TPs, FPs, as well as FNs and consequently precision and recall.

### Development of Novel SV Detection Method

*Dicast* generalizes the concept of consensus calling by implementing a machine learning framework that scores candidate SVs using features derived from genome annotations and sequence alignments. Rather than relying solely on overlapping calls from multiple methods, *dicast* quantifies the probability that each SV represents a true variant by integrating multiple evidence sources. *Dicast* accepts as input a variable number of VCF files containing candidate structural variants calls from existing methods or candidate variants from a population catalog. Input variants are consolidated into a unified format represented by variant type, chromosome, start, end, as well as SV size.

#### Feature Collection

Subsequently, variants are annotated with a range of features describing different aspects of a variant. For that, four bins, two around each breakpoint, are defined around each variant. Let BP_l_ denote the position of the left breakpoint and BP_r_ the position of the right breakpoint. Then the coordinates of the four bins are defined as

- Bin I: [BP_l_ - 52, BP_l_ - 2]
- Bin II: [BP_l_ + 2, BP_l_ + 52]
- Bin III: [BP_r_ - 52, BP_r_ - 2]
- Bin IV: [BP_r_ + 2, BP_r_ + 52]

Since insertions are defined by a single breakpoint, only Bins I and II are considered for this variant type. For each bin, several features derived from the alignment of sequencing reads around the breakpoints are calculated: coverage, insert size, mapping quality, clipped reads, split-reads, and discordant read pairs.

Coverage is assessed by calculating the mean read depth within each bin and normalizing it by the average coverage across the corresponding chromosome. To account for local variation, the standard deviation of coverage within each bin is also computed. In addition to the bins near breakpoints, *dicast* defines four additional bins evenly distributed along the SV body. These internal bins are used exclusively for coverage-based annotations, helping to characterize coverage patterns not only at the breakpoints but across the entire SV. Insert size, defined as the genomic distance between paired-end reads, is measured for each bin by computing the mean insert size of all reads overlapping with a given bin. This mean is normalized by the chromosomal average insert size, and the standard deviation is calculated to capture variability within each bin. Mapping quality is annotated by calculating both the mean and standard deviation of the mapping quality scores of all reads overlapping a given bin. Clipped reads, which are reads that only partially align to the reference genome, can signal potential breakpoints. For each bin, the proportion of clipped reads is calculated as the number of clipped reads divided by the total number of reads overlapping that bin. Split reads, a subset of clipped reads, whose unaligned segments map elsewhere in the genome, are similarly quantified. The proportion of split reads is computed in the same manner as clipped reads. To capture the structural relationship between different regions of a variant, bin connection features are computed. These features quantify the evidence that two distinct bins are connected by abnormal read patterns. For each pairwise combination of bins, two main connection types are assessed:

- Split-read connections: These involve individual reads that are split across two bins, meaning one segment of the read aligns to one bin and the other segment aligns to the second bin. For each bin pair, the proportion of such split reads (relative to all split reads overlapping either bin) is calculated.
- Discordant read pair connections: These involve paired-end reads where one mate aligns to one bin and the other mate aligns to the second bin in an unexpected orientation. For each bin pair, the proportion of these discordant pairs is measured.

In addition to features derived from aligned sequencing reads, *dicast* takes the genomic context of a variant into account. The extensive evaluation of SV detection methods and their underlying technologies has revealed that alignment patterns indicating the presence of a TP structural variant can look vastly different depending on their location in the genome. For example, due to difficult read alignments in repetitive regions, positions of clipped and split reads can be shifted. Therefore, each variant is annotated with a list of features derived from reference genome annotations. These features are described in the Methods section *Evaluation of Technologies*.

#### Training

The collected features serve as the basis to train an *XGBoost* model to distinguish TP SVs from FP artifacts. The set of eight individuals (the ninth one serves as a test set) for which ground truths have been created make up the training set. Variants from unfiltered *VCF* files generated by *delly*, *manta*, *lumpy*, *gridss*, and *cnvnator* were annotated with features as described by the process above. The ground truth datasets generated by the integration of multiple sequencing and one optical mapping technology were used to assign labels (TP or FP) to the unfiltered variants.

#### Evaluation

For deletions and insertions, *dicast* was evaluated on the Freeze 4.0 benchmark set for sample HG002 published by the Human Genome Structural Variation Consortium (HGSVC). Since at the time of writing, no existing benchmark set for duplications was available, the evaluation for this SV type has been performed on the one left-out sample of our newly created ground truth dataset. Both benchmark sets underwent manual curation as described above. Candidate variants were again called using *delly*, *manta*, *lumpy*, *gridss*, and *cnvnator* which were subsequently annotated with the features described above. Additionally, variants from a population catalog published by Ebert et al., 2021, were used as input candidate variants for *dicast*. The population catalog was derived from variants identified in 32 haplotype-resolved human genome assemblies. To avoid data leakage, the test sample HG002 was excluded from the catalog. The trained *dicast* models were used to score variants for each SV type, respectively. A *dicast* score cut-off of 0.4 was used to filter out low-quality, likely FP variants. For the other SV callers, variants which contained the PASS filter tag, were kept. Subsequently, call sets were overlapped with the respective benchmark datasets. Two variants were called overlapping if their breakpoints were located within 500 bp and their size similarity was > 0.7. The number of TP variants per method was determined by the number of benchmark variants overlapping with a called variant. The number of FP was equal to the number of all variants in the callset which did not overlap with a benchmark variant. FN variants were all benchmark variants which did not overlap with any called variant. To determine precision-recall curves, the respective quality scores of all callers were used.

### Evaluation of Novel SV Detection Method in Clinical Scenarios

#### Short- and Long-read Processing

The experimental procedure, alignment, and SV calling of the short-read and long-read data for the three pipelines was performed as previously described for the creation of the ground truth data set with slight adjustments and without assembly-based methods. For the first and third pipeline, we only retained SV calls passing the caller-specific quality filter, i.e., “PASS” variants. We then scored all variants using *dicast* and combined the filtered PacBio and Illumina calls of the same variant type in the third pipeline based on reciprocal overlap with a threshold of 0.7 returning a representative with the highest *dicast* score.

#### Variant Frequency Filtering

We compared our call-sets of each pipeline to four catalogues of common variation (Audano et al., 2019; Ebert et al., 2021; Collins et al., 2020; NCBI nstd186). From each data-set, we discarded rare variants based on allele frequency or similar metrics (0.01 AF for gnomAD-SV v2.1 and NCBI, “Singletons” for the long-read data-sets). Then we clustered the variants identified in our pipelines with the common variants using a reciprocal overlap threshold of 0.7 and only retained those clusters without any known common variation. Given the phenotypic diversity in our limb malformation cases, we expected unique causative variants and filtered out any variants present in multiple individuals, resulting in a set of singletons. We also investigated shared rare variants between patients in a separate analysis not discussed in this publication.

#### Variant Annotation

To identify potential functionally relevant variants, we employed an adapted version of the TADA method (Hertzberg et al., 2022). We used sets of coding and non-coding annotations tailored to the context of limb malformations.

The coding annotations included gene definitions as well as exons, start- and stop-codons, 5’ and 3’ UTRs from ENSEMBL v.110 (Dyer et al., 2025) and developmental-disease associated genes (DECIPHER). We also generated four additional gene sets specific to the patients’ phenotypes : 1) genes correlated with limb development based on a separate scRNA study 2) genes known to be associated with limb development derived from previous work by the Mundlos AG at the Max-Planck-Institute for Molecular Genetics (MPIMG) 3) phenotype associated genes and 4) differentially expressed genes (DEGs). The phenotype associated genes for each patient are based on the patient’s HPO terms (Gargano, et al., 2024). The DEGs are derived from a patient specific RNA-seq analysis.

For non-coding annotations we collected cell-type specific active enhancers determined using cap analysis gene expression (CAGE) from FANTOM5 (Andersson et al., 2014). Given the lack of limb-specific enhancers in this specific resource, we used the entire collection during annotation. We retrieved the entire set of human candidate regulatory elements (cREs) from ENCODE (ENCODE Project Consortium. et al., 2012). While many regulatory elements contained in this set are likely not relevant to limb development, they could indicate regions of potential interest that would warrant further investigation. We also collected a set of experimentally validated enhancers from VISTA (Visel et al., 2007). To reflect the limb development-associated regulatory landscape, we collected a set of cREs identified in mouse embryonic limb tissue (14.5 days - ENCODE ID: ENCFF890IPV) and lifted the data over to GRCh38. Given the lack of publicly available limb-specific CTCF sites in humans, we collected ENCODE ChIP-seq derived CTCF narrow peaks from fibroblast samples (ENCFF882YMD). We also included TAD boundaries derived from our Hi-C analysis both as proxies for windows of increased regulatory interactions and as individual annotations.

Given the set of coding and non-coding annotations, we defined functionally relevant variants as those hitting either limb-developmental associated coding annotations directly or affecting non-coding regulatory elements in the same TAD environment as disease-relevant genes. This process as well as detailed description of data processing and analysis are available in the referenced dissertation (Hertzberg. Diss. 2023). In a final step, we manually curated the functionally relevant variants, resulting in a set of candidate variants for each pipeline. We compared the final candidates of the pipelines using reciprocal overlap with a threshold of 0.7.

#### RNA-seq

The fibroblast samples of the 21 patients were cultured in DMEM with 10% FBS, 1% L-glutamine, and 1% pen-strep. RNA extraction from fibroblasts was performed in all 21 samples using the RNeasy mini kit (Qiagen, Hilden, Germany). The Poly(A) mRNA capture was done using the KAPA mRNA HyperPrep Kit (KR1352 -v5.17) and sequencing was performed on a HiSeq4000 (Illumina) using a single technical replicate (PE75, 50 million fragments per sample). We processed the raw sequencing data with a custom snakemake pipeline: First, we aligned the reads to the GRCh38 reference using *STAR* v.2.7.9a (Dobin et al., 2013). We filtered for reads with a minimum mapping quality (MAPQ) of 5. To identify DEGs for each patient we conducted a one vs. all analysis with respect to the *UCSC hg38 knownGene* library using a custom R script. Briefly, we removed any data mapped to sex chromosomes and counted the reads of each patient per exon with summarizeOverlaps. We then applied the *DESeq2* (v1.26) function DESeq computing the difference between read counts of a single sample and the remaining cohort (Love et al., 2014). Finally, we normalized the results using *vst*, computed the logarithmic Fold-Change (Log2FC) per gene, and returned the top 50 genes with the highest Log2FC as the list of DEGs for each patient.

#### Hi-C

The PCR amplification (4–8 cycles) of the fibroblast samples was done using NEBNext Ultra II Q5 Master Mix (New England BioLabs, M0544). The PCR purification and size selection was carried out using Agencourt AMPure XP beads (Beckman Coulter, A63881). Libraries were sequenced in 75bp, or 100bp paired-end runs on a NovaSeq6000 (Illumina) using between 2 and 5 technical replicates. We conducted the bioinformatic analysis in a custom snakemake pipeline. First, we processed the raw sequencing data using the *Juicer* pipeline (v.1.6) with *bwa* (v0.7.17) for aligning the reads to the GRCh38 reference (Duran et al., 2016). We then merged the deduplicated and filtered reads across technical replicates. For each patient, we applied the pre function from *Juicer-Tools* (v1.22.01) to generate Hi-C maps including all mapped reads with MAPQ⩾ 30. In addition, we merged the deduplicated reads across the entire cohort, creating a high-resolution fibroblast Hi-C map. To determine TAD boundaries, we first generated patient-specific VC SQRT normalized contact maps at 25kb resolution for each chromosome using the Juicer-Tools dump function and then applied *TopDom* (Shin et al., 2016).

#### Rare Disease Cohorts

The experimental procedure of the atrial fibrillation and neuromuscular short-read data were previously described (Hackman et al., 2021; Perrin et al., 2024; Baudic et al., 2024) . The processing, filtering and annotation was done with the same pipeline as for the limb-malformation cases, with adjustments in the prioritization process and annotations given the patients’ phenotypes.

## Acknowledgements

We thank Julien Barc and his team for helpful discussions and assistance in accessing and interpreting the sequencing data published in Baudic et al., 2024.

## Supplementary Material

### Figures

**Sup. Fig. 1:**
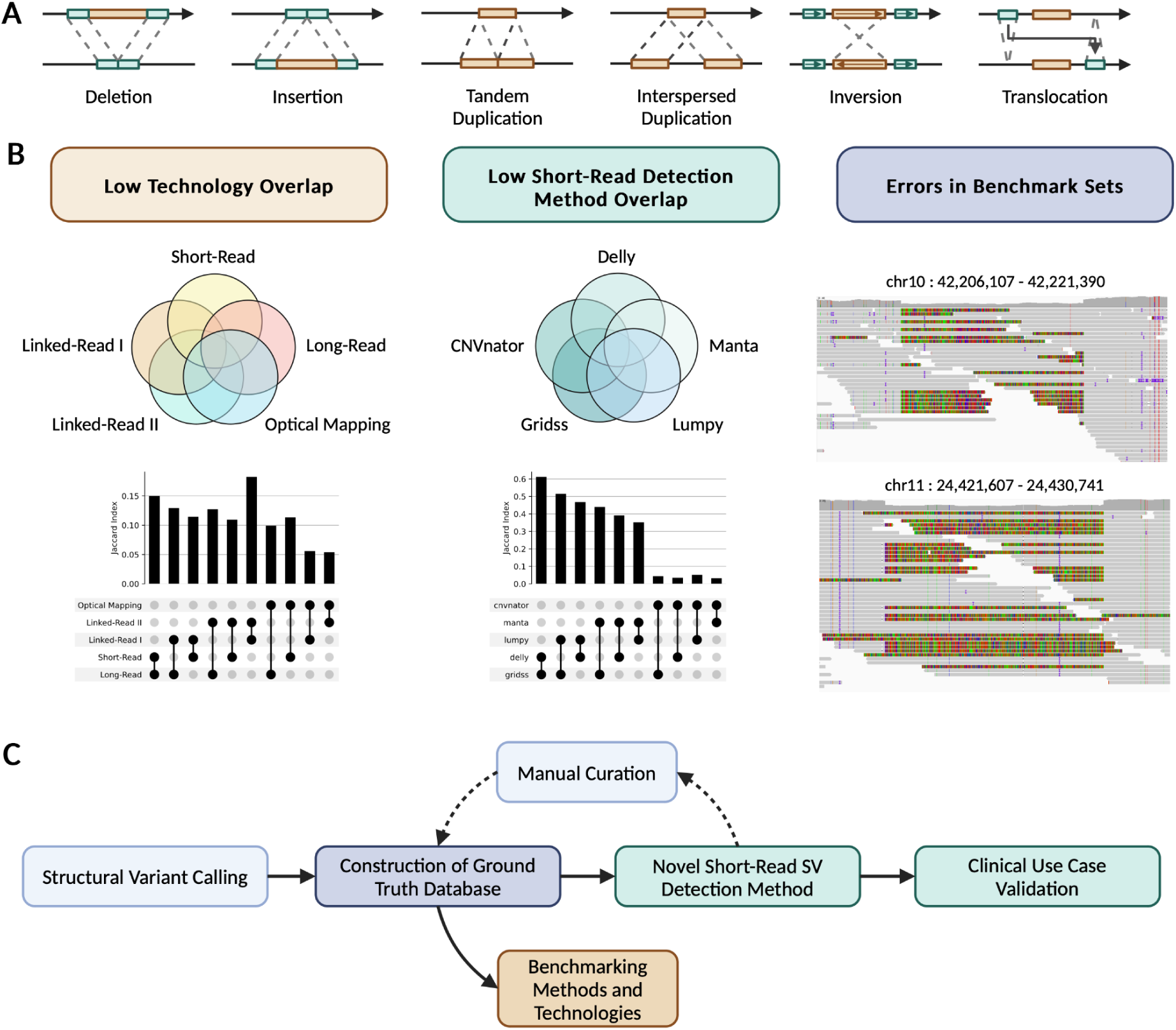
Problem Definition and Workflow Overview. A) Illustration of different types of structural variants. The top arrow represents the reference genome, the bottom arrow represents the patient genome. B) Limitations in structural variant detection. Technology and method overlap are based on one sample of our newly constructed ground truth dataset. *IGV* screenshots depict two example variants in HG002 which show clear alignment patterns corresponding to true heterozygous deletions but which are not part of the HGSVC Freeze 4.0 benchmark set. C) Overview of the performed steps in the paper.

**Sup. Fig. 2:**
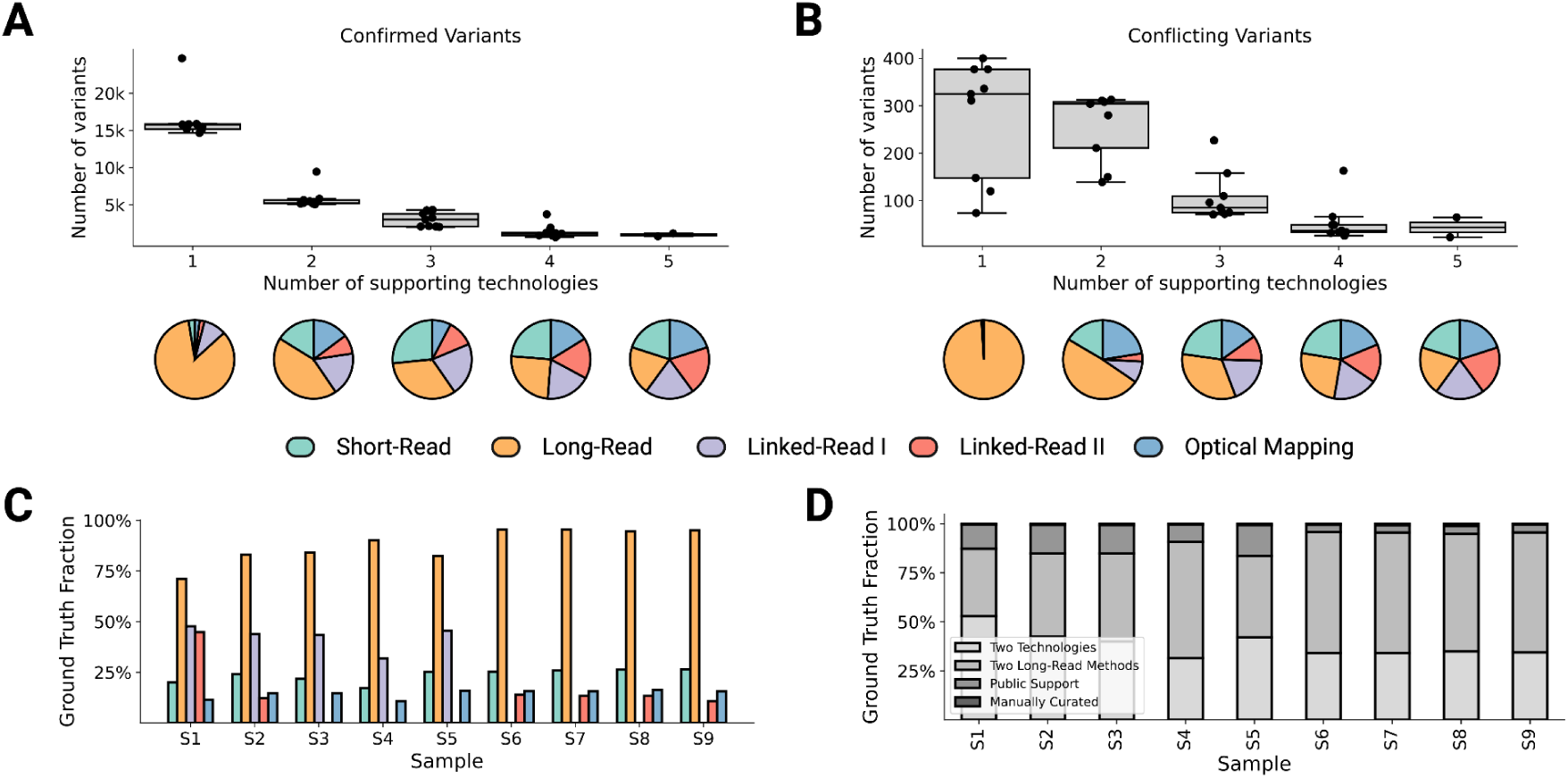
Details on the ground truth SV dataset. A) Number of confirmed variants per number of supporting technologies. Each dot represents one sample in the benchmark dataset. The pie charts beneath each number of supporting technologies show the distribution of variants detected by each technology. B) Number of conflicting variants per number of supporting technologies. C) Fraction of ground truth variants supported by each technology. D) Fraction of ground truth variants per confirmation method. Used strategies to confirm a variant were: Two technology support, two long-read method support, present in public database, manual curation.

**Sup. Fig. 3.**
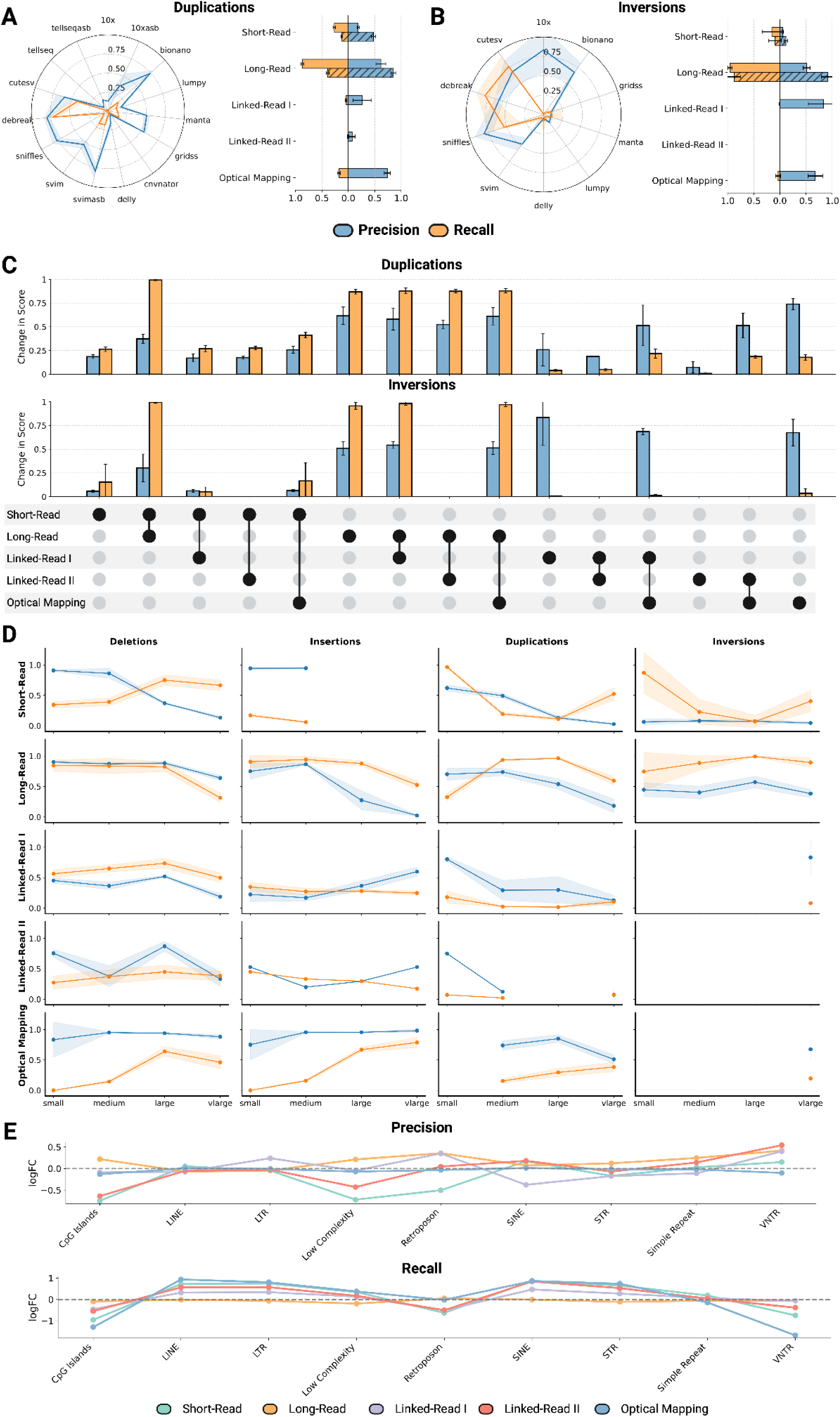
Comparative analysis of methods and sequencing technologies for structural variant detection. Precision and recall for duplication detection. The blue line represents precision, while the orange line represents recall. The confidence interval indicates the standard deviation for precision and recall across nine samples. The horizontal bar plots compare the precision and recall when combining multiple methods within one technology. Striped bars describe a more than 2/n caller strategy, while non-striped bars describe a union strategy. (Short-Read: Illumina, Long-Read: PacBio CLR, Linked-Read I: 10x Genomics, Linked-Read II: TELL-Seq, Optical Mapping: Bionano) B) Precision and recall for inversion detection. C) Evaluation of combining multiple sequencing technologies for SV detection. Multiple methods within one technology were combined using the union strategy. Error bars indicate standard deviation across all samples. D) Precision (blue) and recall (orange) across technologies, SV types, and size categories. Multiple methods within one technology were combined using the union strategy. Shaded areas reflect the standard deviation across nine samples. E) Impact of genomic context on SV detection performance. Depicted is the log2 fold change of precision and recall within a particular genomic context compared to the genome-wide precision and recall.

**Sup. Fig. 4:**
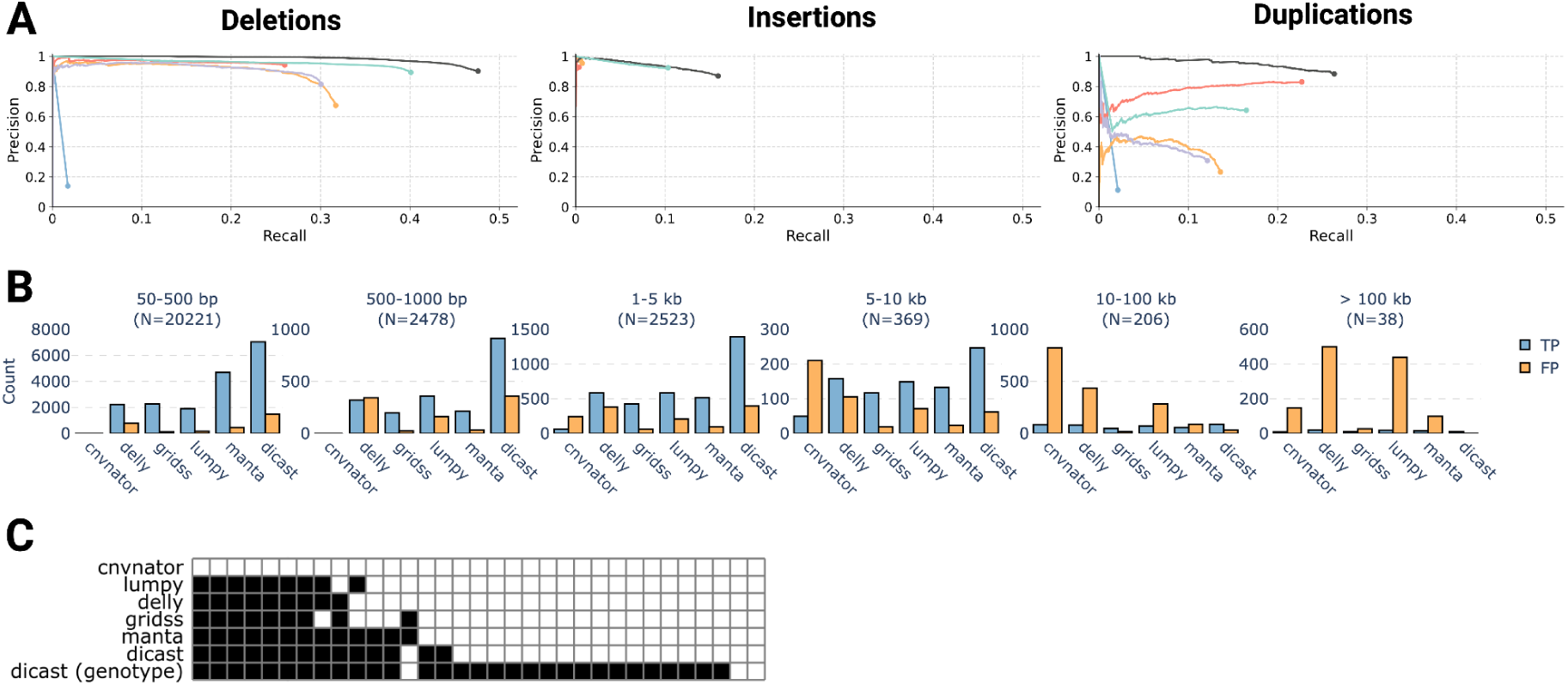
Novel machine learning tool reliably distinguishes true variants from artifacts. A) Precision-recall curves for *dicast* when only receiving the unfiltered callsets from the five SRS-based callers as input candidate variants and not an additional population catalog. B) Number of TP and FP variants across different SRS-based detection methods and size categories. *N* depicts the number of TP variants in the respective benchmark dataset. Considered are deletions, insertions, and duplications. C) Detection performance for deletions within challenging medically relevant genes (CMRG) as defined by Wagner et al., 2022. Each square represents one deletion. Filled squares indicate whether a method detected a variant.

**Sup. Fig. 5:**
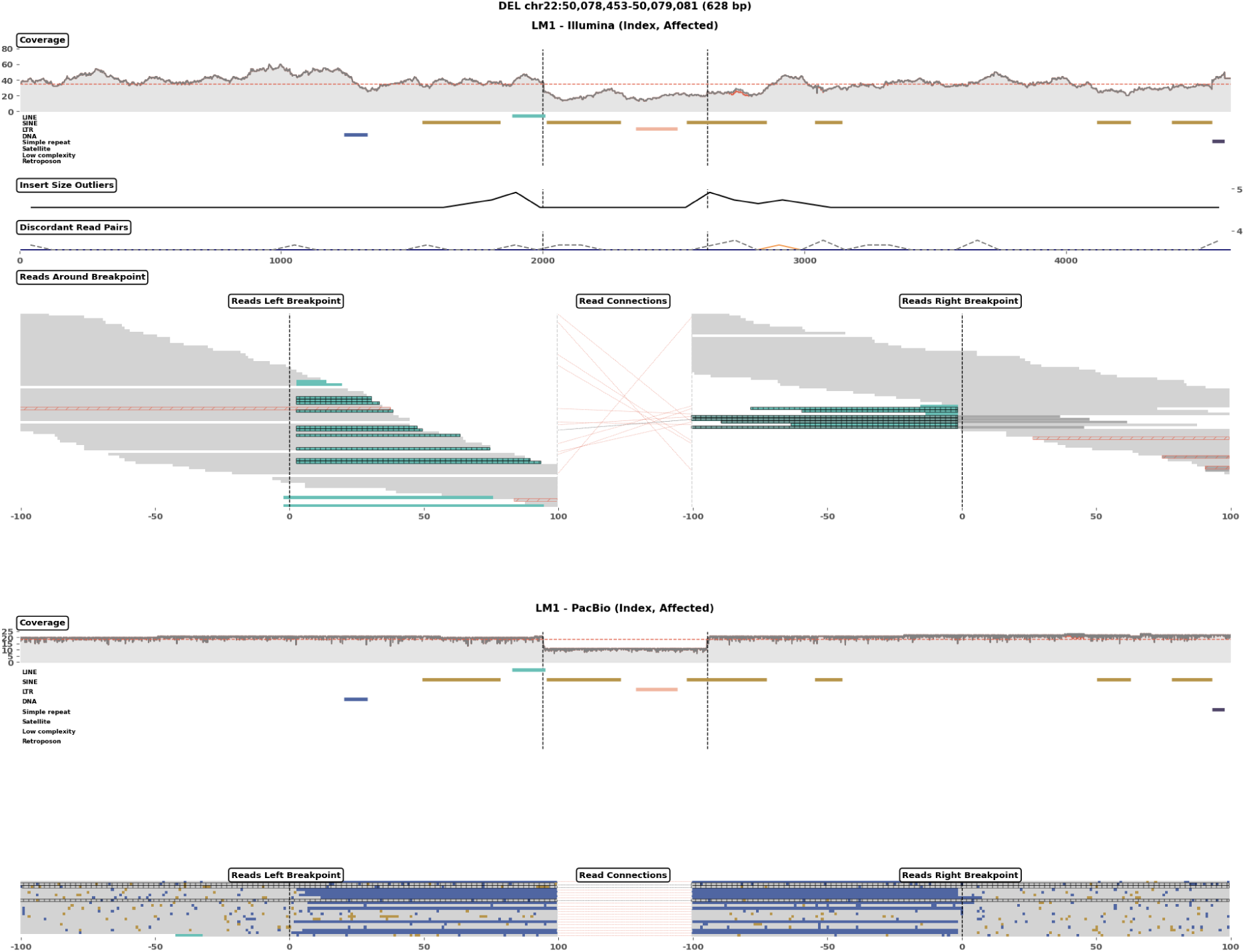
Manually confirmed calls with high dicast prediction (0.998) and clear evidence in both short- and long-read sequencing. Visualizations were created with cuban (See Sup. Fig. 12).

**Sup. Fig. 6:**
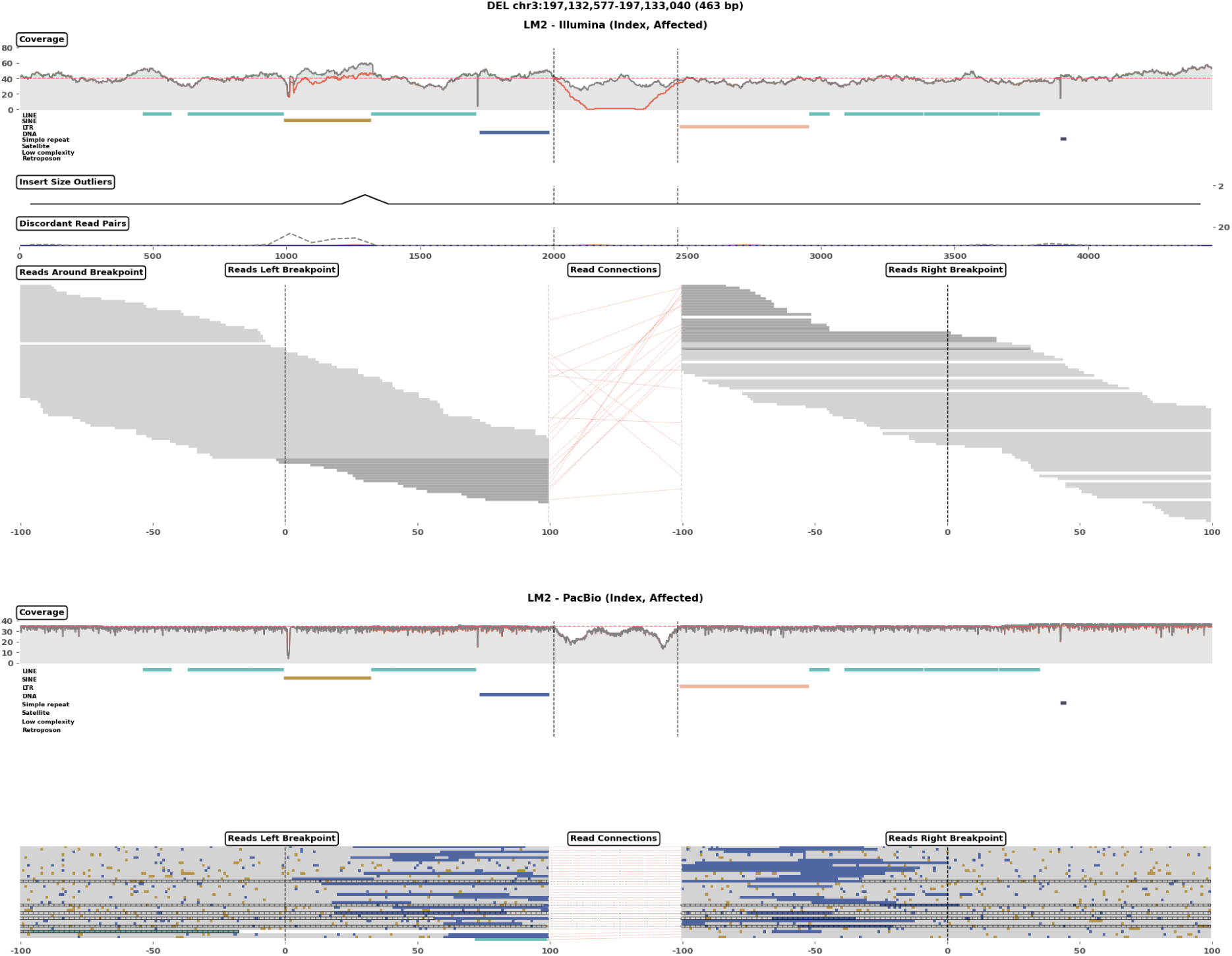
Manually confirmed call with medium Dicast score (0.572) with conflicting evidence short- and long-read evidence. Visualizations were created with cuban (See Sup. Fig. 12).

**Sup. Fig. 7:**
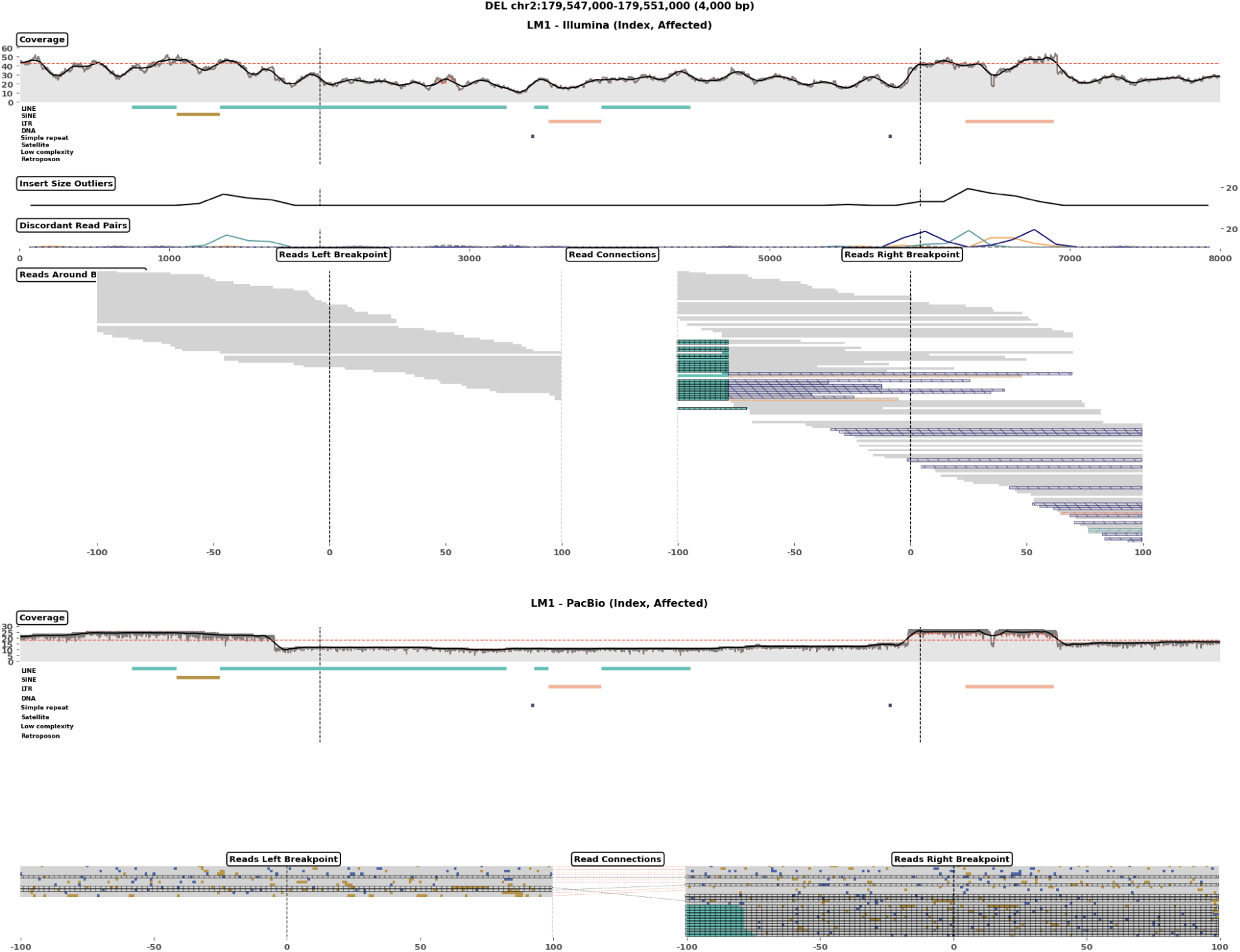
Manually confirmed call with low Dicast score (0.1399) with inaccurate breakpoints. Visualizations were created with cuban (See Sup. Fig. 12).

**Sup. Fig. 8:**
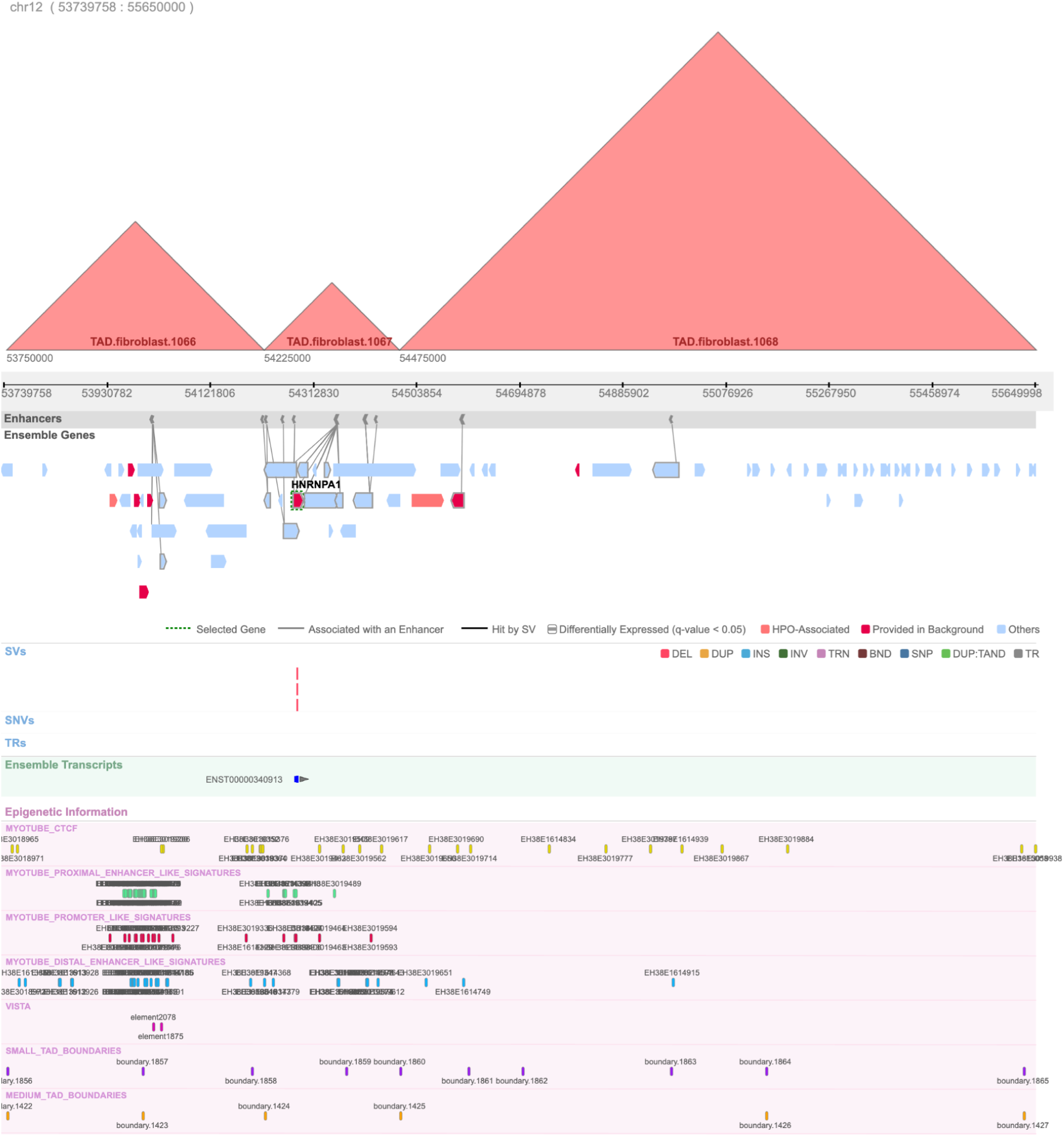
Visualization of the regulatory region affected by the NMD01-NMD03 deletions generated with the Lucid Genome Suite. The figure shows several tracks of relevant coding and non-coding annotations starting with TAD annotations derived from our fibroblast Hi-C analysis and Enhancer signatures from ENCODE connected to the ENSEMBL gene they most likely regulate through the Hi-C interaction frequencies. The following tracks show the variants (SVs, SNVs and TRs) in all samples of the NMD cohort. The canonical transcript of the gene in focus (HNRNPA1) is shown in the next track. Finally, several additional regulatory tracks are included: CTCF binding sites, enhancer and promotor signatures derived through ChIP-seq experiments conducted on myotube samples (ENCODE), VISTA enhancers and TAD boundaries called with various resolutions.

**Sup. Fig. 9:**
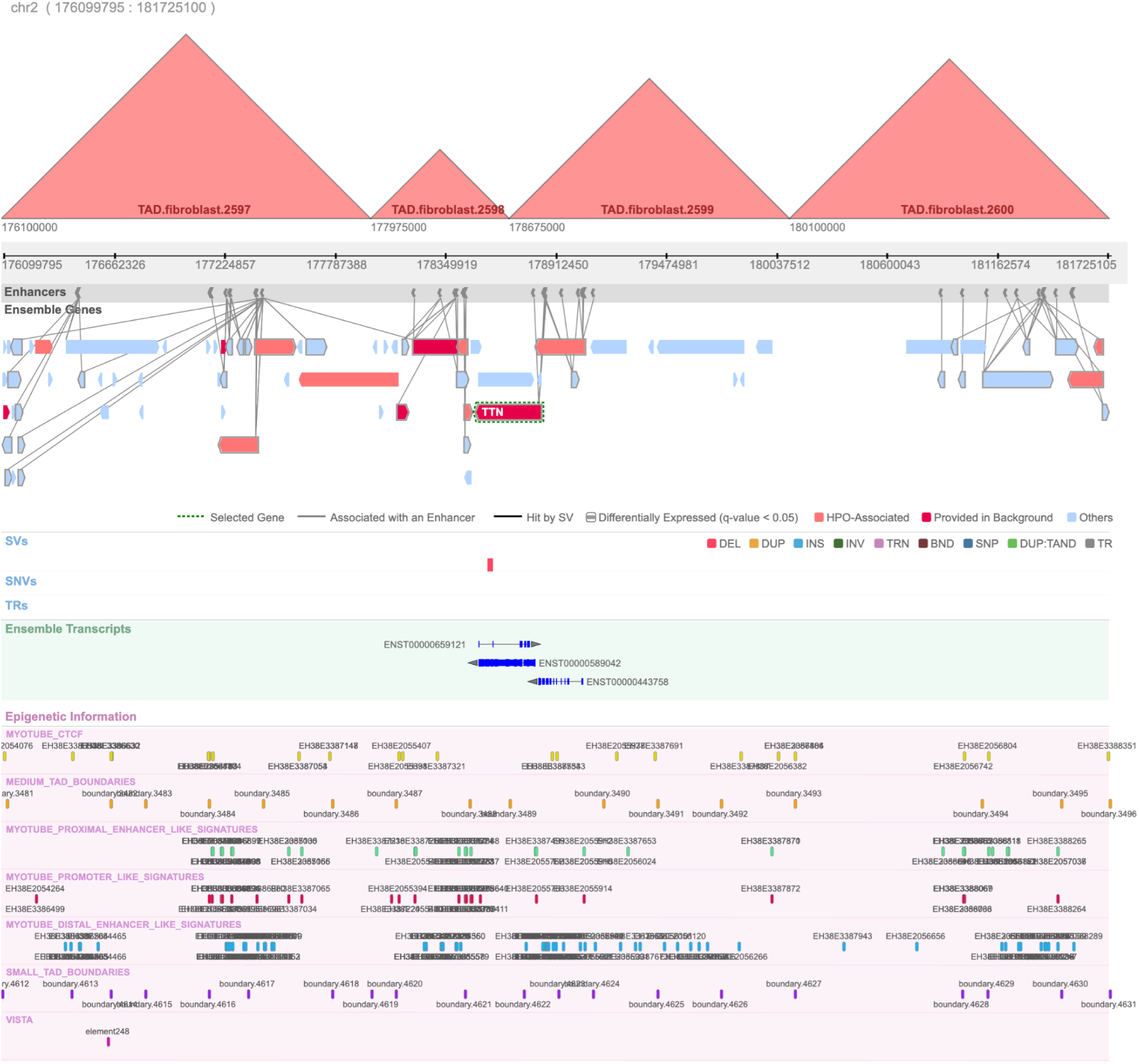
Visualization of the regulatory region affected by the NMD04 deletion generated with the Lucid Genome Suite.

**Sup. Fig. 10:**
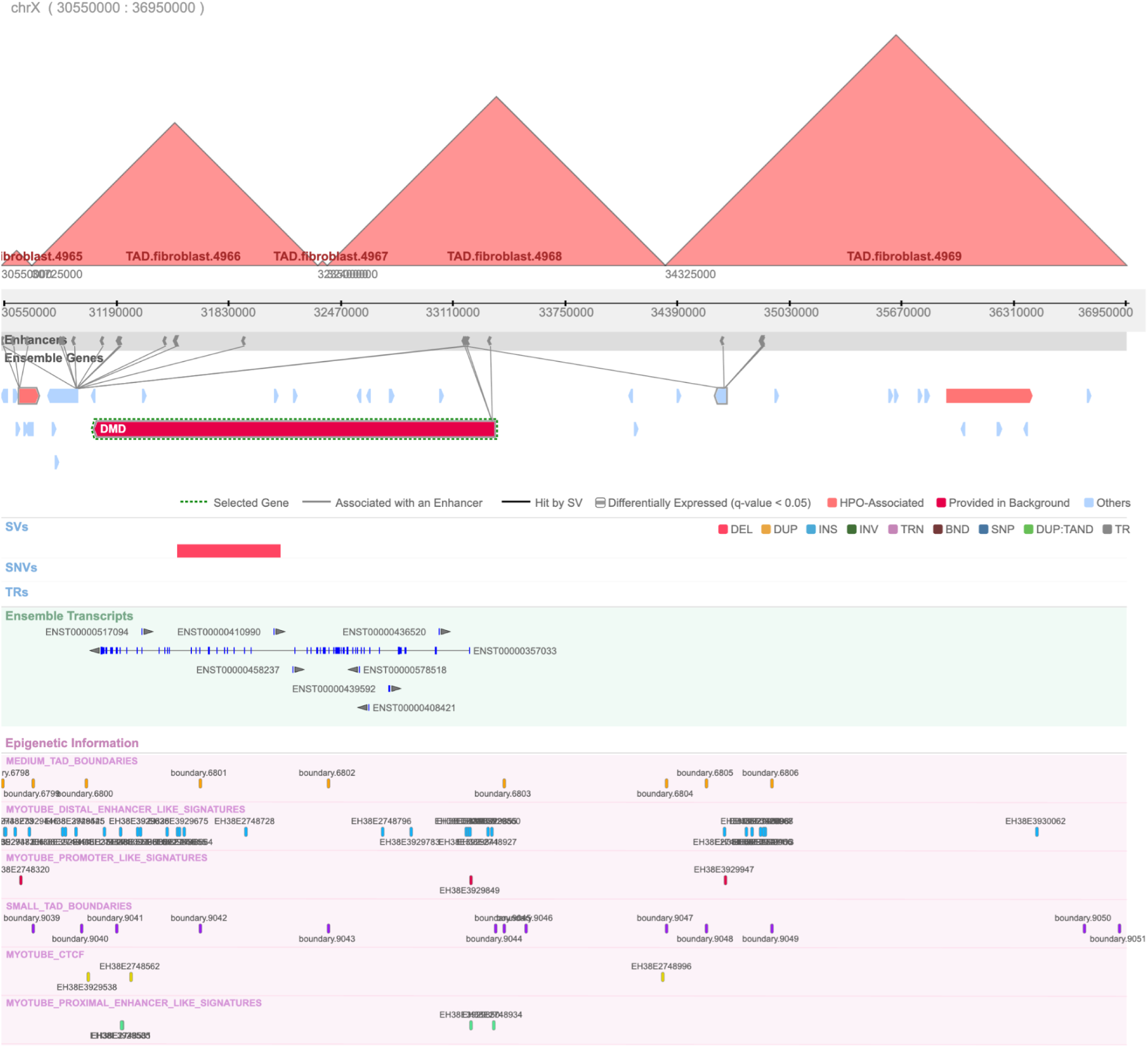
Visualization of the regulatory region affected by the NMD05 deletion generated with the Lucid Genome Suite.

**Sup. Fig. 11:**
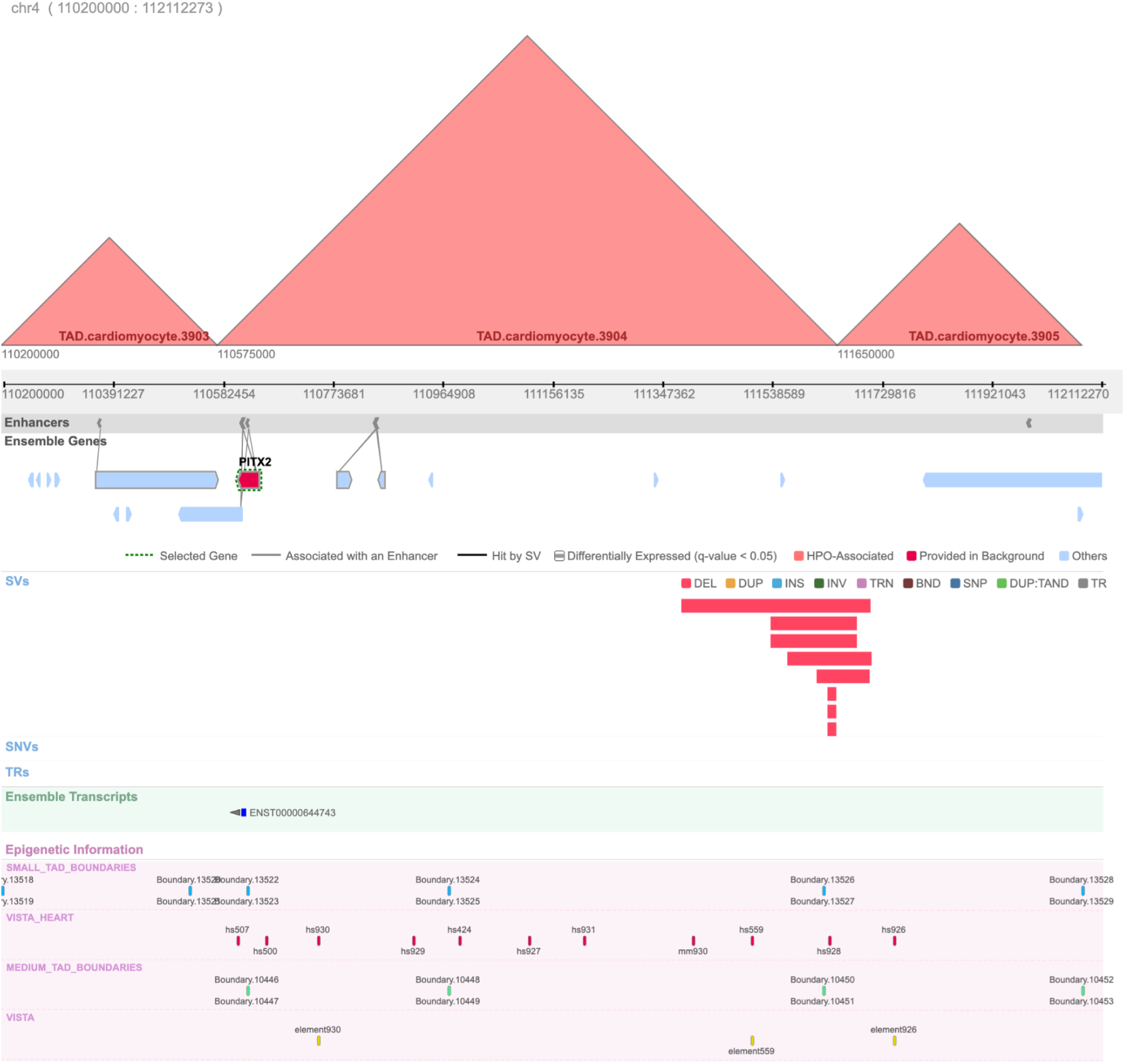
Visualization of the regulatory region affected by the AF deletions generated with the Lucid Genome Suite. The tracks include TADs derived from cardiomyocytes, enhancer signatures from the same cell type connected through Hi-C interaction frequencies to the genes they most likely regulate. In the last part of the figure all VISTA enhancers and those specifically active in heart cell-types in combination with TAD boundaries of various resolutions are shown. The remaining tracks are described in Sup. Fig S8.

**Sup. Fig. 12:**
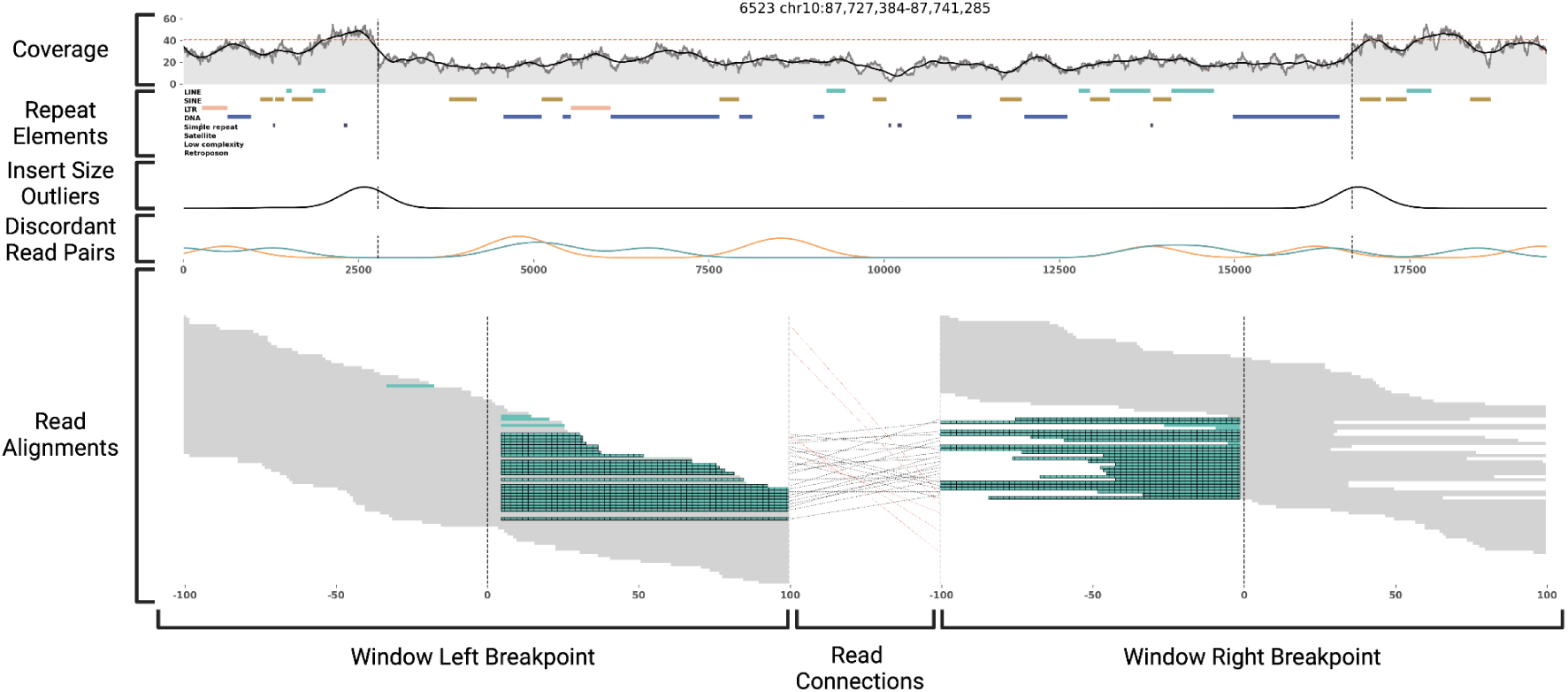
Architecture of cuban plots. Top tracks depict coverage, repetitive elements insert size outliers and discordant read pairs across the whole length of a given variant. The dotted red line within the coverage track indicates the chromosomal average coverage. Discordant read pairs include pairs in reverse-forward (dark blue), forward-forward (green) and reverse-reverse (orange) orientation. The bottom segment visualized CIGAR strings in two 200bp windows around each breakpoint. Light gray indicates a match/mismatch between the read and the reference, while green indicates soft-clipped bases. A black grid overlay indicates a split-read. Connections between reads around each breakpoint are shown between the two windows. Black lines indicate two segments of a split-read, while red lines connect two mates of the same read pair.

### Tables

**Sup. Table S1:** Known pathogenic SVs used in the clinical diagnostic evaluation of *dicast*.

| Sample | Type | Chr | Start | End | Size | Dicast Score | Regulatory Region | Reference / DOI |
| --- | --- | --- | --- | --- | --- | --- | --- | --- |
| NMD01 | DEL | 4 | 54284192 | 54284353 | 161 | 0.991 | Figure S8 | 10.1212/NXG.00000000000000632 |
| NMD02 | DEL | 4 | 54284192 | 54284353 | 161 | 0.592 | Figure S8 | 10.1212/NXG.00000000000000632 |
| NMD03 | DEL | 4 | 54284192 | 54284353 | 161 | 0.994 | Figure S8 | 10.1212/NXG.00000000000000632 |
| NMD04 | DEL | 2 | 178567644 | 178597451 | 29807 | 0.927 | Figure S9 | 10.1136/jmg-2023-109473 |
| NMD05 | DEL | X | 31551000 | 32135379 | 584379 | 0.965 | Figure S10 | Manuscript in Preparation |
| AF01 | DEL | 4 | 111633898 | 111649216 | 15318 | 0.99 | Figure S11 | 10.1038/s41467-024-47739-x |
| AF02 | DEL | 4 | 111633898 | 111649216 | 15318 | 0.954 | Figure S11 | 10.1038/s41467-024-47739-x |
| AF03 | DEL | 4 | 111633898 | 111649216 | 15318 | 0.995 | Figure S11 | 10.1038/s41467-024-47739-x |
| AF04 | DEL | 4 | 111615000 | 111707169 | 92169 | 0.465 | Figure S11 | 10.1038/s41467-024-47739-x |
| AF05 | DEL | 4 | 111534000 | 111685120 | 151120 | 0.947 | Figure S11 | 10.1038/s41467-024-47739-x |
| AF06 | DEL | 4 | 111534000 | 111685120 | 151120 | 0.984 | Figure S11 | 10.1038/s41467-024-47739-x |
| AF07 | DEL | 4 | 111563720 | 111710000 | 146280 | 0.909 | Figure S11 | 10.1038/s41467-024-47739-x |
| AF08 | DEL | 4 | 111380000 | 111708000 | 328000 | 0.63 | Figure S11 | 10.1038/s41467-024-47739-x |

